# Transcriptional plasticity of ventral tegmental area neurons induced after cessation of chronic cocaine exposure is required for incubation of cocaine seeking

**DOI:** 10.64898/2026.06.30.735717

**Authors:** Arthy Narayanan, Kendyl N. Laumann, Aaron T. Halvorsen, Sasha Burwell, Andrew I. Aldridge, Jinjutha Cheepluesak, Xiaolin Wei, Yarui Diao, Christie D. Fowler, Anne E. West

**Author notes:** To whom correspondence should be addressed: Anne E. West, Christie D. Fowler.

## Abstract

Chronic cocaine triggers changes in brain function that persist long after drug taking has ceased. Relapse to substance use during abstinence can be triggered by drug-associated cues, implicating the drug-free period after chronic cocaine as a time when the probability of relapse is modulated by plasticity mechanisms. Here, we studied the regulation and function of transcriptional plasticity activated in a time-dependent manner in dopaminergic and glutamatergic neurons of the mouse ventral tegmental area (VTA) over a drug-free period after chronic cocaine. We used in vivo dCas9/CRISPR inhibition to demonstrate a causal role for Brain-Derived Neurotrophic Factor transcription in the incubation of cocaine seeking during abstinence, and we used single-nucleus sequencing to identify a program of gene regulation induced in VTA neurons selectively by the prolonged absence of cocaine. These data advance the understanding of plasticity mechanisms that modulate reward functions of VTA neurons, which may be a main factor propagating relapse in substance use disorders.

## Introduction

Chronic exposure to drugs of abuse, such as the psychostimulant cocaine, have long-lasting consequences on brain function even after drug consumption has ceased. Although the neural underpinnings of these memory traces remain largely unknown, it is thought that drug-initiated changes in neural function continue to evolve during periods of drug abstinence driving the high likelihood of cue-induced relapse (Gawin & Kleber, 1986; O’Brien et al., 1992). Thus, discovering the precise molecular mechanisms of neural plasticity that are active during drug-free periods after chronic exposure is important to identify innovative interventions that can disrupt the cycle of substance use, abstinence, and relapse in addiction.

Drugs of abuse act on the reward circuitry in the brain, which comprises reciprocal connections between dopamine (DA) neurons of the ventral tegmental area (VTA) with neurons of the nucleus accumbens, hippocampus, and prefrontal cortex (Russo & Nestler, 2013). The VTA is of particular interest due to the centrality of this brain region in reward-related and goal-directed behaviors (Morales & Margolis, 2017). The neurobiology of relapse has been studied in rodents using the incubation of drug seeking paradigm (Chow et al., 2025; Pickens et al., 2011), in which time-dependent increases in lever pressing during drug abstinence serve as a measure of incubation that is proposed to arise from the plasticity of brain networks that underlie drug seeking. Prior studies using the incubation of cocaine seeking model identified an accumulation of Brain Derived Neurotrophic Factor (BDNF) in the VTA that temporally tracks the increase in cocaine seeking behaviors during abstinence (Grimm et al., 2003). BDNF is a secreted neurotrophic factor with well-established roles in synaptic plasticity. Upon withdrawal from repeated cocaine, excitatory synapses onto VTA DA neurons become primed for potentiation in a manner that is dependent on signaling through BDNF and its receptor TrkB (Pu, Liu et al. 2006). Furthermore, injecting BDNF protein directly into the VTA during the early part of the abstinence period is sufficient to both immediately and persistently enhance cocaine seeking behavior (Lu et al., 2004).

These data raise the possibility that VTA BDNF could play a role in the long-term molecular and circuit mechanisms of drug craving that arise during abstinence from cocaine. However, a number of key questions remain unanswered. First, the VTA is comprised of multiple cell types including several different kinds of neurons with distinct projection patterns and behavioral functions (Morales & Margolis, 2017). Single-cell sequencing studies have shown that *Bdnf* mRNA is found in glutamatergic as well as dopaminergic neurons of the VTA (Phillips et al., 2022), however which cell types are responsible for the increased BDNF observed in the VTA during abstinence is unknown. Second, although cocaine is known to activate DA-regulated signaling pathways that can promote gene expression via modification of transcription factors and chromatin regulators, the mechanisms by which abstinence from cocaine induces slowly evolving and persistent changes in gene expression is poorly understood. One study reported that VTA induction of BDNF after cocaine withdrawal is associated with specific histone changes at promoter I of the *Bdnf* gene (Schmidt et al., 2012), however whether activation of this promoter is required either for *Bdnf* induction during abstinence or for the incubation of drug seeking after cocaine remains to be tested. Finally, it is likely that additional genes are regulated in VTA neurons that may contribute to the incubation of cocaine craving. For example, one study found that histone H3 in the VTA of rats is subject to progressive dopaminylation at glutamine 5 (H3Q5dop) after extended cocaine self-administration (Lepack et al., 2020), however whether there are coordinated programs of VTA gene expression during periods of drug abstinence remains poorly understood.

Here, to address these questions, we performed multiplex single-molecule fluorescent in situ hybridization (smFISH) to quantify *Bdnf* expression in VTA cell types across time during withdrawal from repeated cocaine. We developed dCas9/CRISPR interference tools to first functionally validate the genome regulatory elements that mediate *Bdnf* transcription during abstinence and then to demonstrate that transcriptional induction of *Bdnf* in the VTA in vivo is necessary for the increase in seeking behavior during cocaine abstinence. Finally, we used genome-wide single nucleus RNA sequencing of neurons from the VTA to identify a program of gene expression induced during abstinence after repeated cocaine and we generated genome-wide chromatin conformation maps from VTA DA neurons to identify the regulatory elements and transcription factors that orchestrate this transcriptional program. These data advance understanding of the plasticity mechanisms that operate in DA neurons of the mouse VTA during periods of abstinence after chronic cocaine exposure and that may modulate the likelihood of relapse to drug seeking.

## Materials and Methods

### Mice

Rosa-LSL-dCas9-KRAB mice (#033066, RRID:IMSR_JAX:033066) (Gemberling et al., 2021), *Gt(ROSA)26Sor^tm5(CAG-Sun1/sfGFP)Nat^* (INTACT) mice (#021039, RRID:IMSR_JAX:021039) (Mo et al., 2015), and *Slc6a3^tm1.1(cre)Bkmn^*(DAT^IRES*cre*^) mice (#006660, RRID:IMSR_JAX:006660) (Bäckman et al., 2006) were purchased from the Jackson Laboratory and bred in our vivarium. Timed-pregnant CD1 IGS mice were purchased from Charles River Laboratories (Strain 022, RRID:IMSR_CRL:022) and used for all neuronal culture experiments. All experiments were conducted in accordance with animal protocols approved by the Duke University Institutional Animal Care and Use Committee and UC Irvine Institutional Animal Care and Use Committee. Both male and female mice were used for all experiments in this study.

### RNA fluorescent in situ hybridization (FISH)

Mice were anesthetized with isoflurane then freshly harvested brains were flash frozen in isopentane chilled in an ethanol and dry ice bath prior to embedding in OCT. Brains were stored at –80⁰C prior to sectioning. Brains were cryosectioned at 20µm thickness and coronal sections mounted on Super Frost Plus slides. To select brain slices for the VTA, immunocytochemistry for tyrosine hydroxylase (TH) was performed on three midbrain slices using a rabbit anti-TH antibody (Pel-Freez Biologicals Cat# P40101-150, RRID:AB_2617184, Dilution 1:1000). The slice with the strongest TH signal and flanking slices were selected for FISH, typically at AP =-3.27mm. We performed RNAScope for FISH following the protocol from the Advanced Cell Diagnostics (ACD Bio) RNAScope Multiplex Fluorescent Assay v2 kit (Cat no. 323110). We used the following probes (Advanced Cell Diagnostics, Newark, CA): Mm-*Bdnf*-C2 (Cat no. 316031-C2), Mm-*Slc6a3*-C1 (Cat no. 315441-C1), Mm-*Slc17a6*-C3 (Cat no. 319171-C3). Slides were counterstained with DAPI to identify nuclei and coverslipped using ProLong Gold mounting medium. The RNAscope signal was imaged at 40X on a Leica SP8 confocal microscope and. A total of 4 images were captured per slice per animal (2 medial VTA images and 2 lateral VTA images).

### RNA-FISH Image Analysis

The images were analyzed and the *Bdnf* signal was quantified using Fiji/ImageJ 1.54p (RRID:SCR_003070). Eight 0.5µm z-steps centered on the largest diameter DAPI signal were collapsed into a sum projection for each cell. Background fluorescence for the *Bdnf* channel was calculated from 4 sample ROIs within each image and subtracted from the image. To determine the threshold signal required for VTA cell type classification, cells exhibiting very low—but clearly detectable—signal for each cell type-specific probe (*Slc6a3* or *Slc17a6*) were first identified. The integrated density of these cells was measured and used to establish the threshold for each respective probe. Only if a cell had a *Slc6a3* integrated density higher than the threshold was it considered at DAT cell. Similarly, only if a cell had a *Slc17a6* integrated density higher than the threshold was it considered at VGLUT2 cell. 50-70 cells were sampled per image. An ROI was drawn around the DAPI signal per cell, and the integrated density for each probe was measured. The thresholding allowed for classification of specific VTA cell types, within which *Bdnf* signal was measured. Final *Bdnf* signal was calculated by subtracting the background *Bdnf* fluorescence from the measured fluorescence within each cell.

### Non-contingent cocaine administration

Mice were intraperitoneally (i.p.) injected with saline as a vehicle control or 20mg/kg cocaine (Sigma Cat no. C5776) for seven consecutive days in the homecage. One day (abstinence day 1, AD1) or 14 days (AD14) after treatment was stopped, the mice were deeply anesthetized with isoflurane before rapid decapitation. For the dCas9-KRAB mice, the brains were processed for RNA FISH. For the DAT-Cre x INTACT mice, the VTA was microdissected and flash frozen in isopentane chilled in an ethanol/dry ice bath, following which the samples were processed for nuclear isolation.

### Viruses

For viral infection of neuron cultures, we used pLV-hUbC-dCas9 KRAB-T2A-GFP plasmid (Addgene #67620, RRID:Addgene_67620) and FUGW-U6-gRNA (Addgene #14883, RRID:Addgene_14883). These constructs were packaged as lentiviruses using HEK293T cells with VSVg and d8.9 plasmids following standard procedures, purified by ultracentrifugation, and functionally titered in both HEK293T cells and cultured cortical neurons using imaging for GFP. The gRNA sequences provided in **Table S1** were designed using online sgRNA designing software (E-CRISP, IDT, GT-Scan). The sequences were cloned into the BsmBI restriction site of FUGW-U6-gRNA. gRNAs that demonstrated strong repression of specific *Bdnf* transcripts in culture were selected for *in vivo* experiments. For viral infection of neurons *in vivo* in the dCas9-KRAB transgenic mice, the gRNA sequences were cloned into the SapI restriction site in the AAV plasmid U6-sgRNA-hSyn-Cre-2A-EGFP-KASH, which allows for co-expression of the gRNA, Cre recombinase, and GFP in infected neuron (Addgene #60231, RRID:Addgene_60231) and packaged into AAVs in serotype Rh10 by the Duke Viral Vector Core. The gRNAs used in the *in vivo* experiments to target *Bdnf* regulatory elements were P1g3, P4g3, and IEg3.

### Mouse embryonic cortical neuron cultures

Neuron cultures were generated using male and female embryonic mouse cortices (E16.5 CD1) as previously described (McDowell et al., 2010; Tao et al., 2002). Briefly, the cortices were dissected and digested with papain (Worthington, Cat no. LS003126). The tissue was triturated by repetitive pipetting and cells were counted. Neurons were plated on PDL coated plates in 10% serum-containing media, then 4 hours later the media was changed to Neurobasal + B27 (Gibco, Cat no. 21103-049, Cat no. 17504-044). Neurons were infected (MOI = 1) on day in vitro 1 (DIV1) for 6 hours with lentivirus in Basal Media Eagle (Sigma Aldrich) with 0.4mg/ml added polybrene, then cells were returned to conditioned media. Isotonic membrane depolarization using 55mM extracellular KCl (Tao et al., 1998) was performed on DIV7 for 6 hours to induce *Bdnf* transcription.

### Reverse transcription and quantitative PCR

RNA was harvested using TRIzol (Invitrogen) and treated with DNAse I (New England Biolabs, Cat No. M0303S) prior to cDNA synthesis using Superscript II (Invitrogen, Cat No. 18064014) and Random Hexamers (ThermoFisher Scientific, Cat No. SO142). Quantitative SyBR green PCR was performed on a QuantStudio^TM^ 3 Real-Time PCR Instrument (Applied Biosystems, A28132) using intron-spanning primers (**Table S2**).

### Stereotaxic VTA injections for virus validation

All surgeries were conducted under aseptic conditions, and body temperature was maintained at approximately 36°C with a heating pad. Mice (7-14 weeks old) were anesthetized with 2-3% isoflurane and placed on the stereotaxic apparatus (Kopf Instruments, 1900). Buprenorphine 0.5 mg/ml (extended-release, 0.5mg/kg) was administered subcutaneously, followed by the application of lubricating ophthalmic ointment to protect the corneas from drying out. The fur on the head was removed, and a midline incision was made to expose the skull. A few drops of bupivacaine 0.25% were administered around the incision as a topical analgesic. The skull was cleaned with the application of 3% hydrogen peroxide and leveled across both the anterior-posterior and medial-lateral axes. Cranial burr holes were made with a carbide drill bit (R1001/4c) at AP =-3.2 mm and ML = ±0.5 mm. For viral targeting of the VTA, the virus used was at a final titer of 5E+12. The virus was loaded in the glass pipette connected to the microinjector (Narishige, MO-10). The pipette tip was lowered to DV =-5.0 and 4.5 mm, and the injections happened at a rate of ∼100 nl/minute for a total volume of 100 nl per site, followed by a 10-minute wait to allow for diffusion of the virus before withdrawing the pipette. The burr holes were then covered with bone wax, and the incision was sutured shut, and covered with a Neosporin antibiotic ointment. The mice received a subcutaneous injection of hydration fluid (sterile saline with 5% dextrose, 1.5ml) and were monitored for 48 hours post-surgery with access to DietGel recovery supplement (ClearH2O, 72-06-5022).

### Contingent cocaine abstinence paradigm

For behavioral studies, subjects were tested during the dark phase of the reverse 12:12 light:dark cycle and food restricted to ∼90% of free feeding weights. The dCas9-KRAB male and female mice (at least 6 weeks of age) were trained to press levers in the operant chamber (Med Associates) for chow food pellets (20 mg; TestDiet) up to a fixed ratio 5, time out 20 sec schedule of reinforcement (FR5TO20) in 1 hr daily sessions. Each session was performed using 2 retractable levers (one active, one inactive) that extend 1 cm into the chamber. Completion of the response criteria on the active lever resulted in the delivery of a food pellet. Data were automatically recorded by the Med Associates software. Mice were then prepared for surgical implantation of the jugular intravenous catheter. *Intravenous surgery*: Mice were pretreated with meloxicam (0.2 mg/kg, oral), shaved at the neck and back, and then anesthetized with an isoflurane (1%–3%)/oxygen vapor mixture in preparation for surgery under aseptic conditions. Catheters comprised of a 6 cm length of silastic tubing fitted onto a guide cannula bent at a curved right angle and encased in dental acrylic. Catheter tubing was subcutaneously passed from the animal’s back into the right jugular vein. A 1 cm length of the catheter tip was inserted into the vein and secured with surgical silk suture. Incision sites were closed with 4-0 surgical sutures. Thereafter, animals were provided at least 72 hrs to recover from surgery, during which time the analgesic meloxicam was administered. *Cocaine self-administration*: After recovery, mice were permitted to respond for cocaine infusions (0.3 mg/kg/infusion) under the FR5TO20 schedule of reinforcement. Reward infusions delivered 0.0367 mL of intravenous cocaine over 3 seconds by a Razel syringe pump (Med Associates) and were paired with a small LED cue light located directly above the active lever during the 20 sec time out. Inactive lever responses were recorded but were noncontingent, and data were automatically recorded by the Med Associates software. Subjects intravenously self-administered cocaine in 1 hr daily sessions across 10 sessions to allow for chronic exposure conditions.Incubation of cocaine craving, day 1: The day after the last cocaine self-administration session, the mice underwent incubation testing (abstinence day 1). The incubation test was conducted by the mice being placed in the same operant chambers to lever press under the FR5TO20 schedule, but completion of the response criteria only resulted in activation of the cue light, without any cocaine infusions. Immediately following the first incubation test session, half of each group (cocaine or saline) were randomly assigned to receive stereotaxic microinjection of the virus with gRNA. *Stereotaxic injections*: Subjects were anesthetized with 2-3% isoflurane, meloxicam (0.2 mg/kg, oral) was administered, and the head was shaved and cleaned, followed by the application of lubricating ophthalmic ointment. Mice were placed on the stereotaxic apparatus (Kopf Instruments) under aseptic conditions, lidocaine was injected below the skin, and then a midline incision was made to expose the skull. The skull was cleaned and leveled across both the anterior-posterior and medial-lateral axes. Cranial burr holes were made with a carbide drill at AP =-2.9 mm and ML = ±0.5mm, bilateral. The virus was loaded in a stainless-steel injector connected to a microinjector syringe (Hamilton). The pipette tip was lowered to DV = - 4.25 mm, and the injection was administered at a rate of 100 nl/minute for a total volume of 200 nl per side, followed by a 7-minute wait to allow for diffusion of the virus. The incision was closed shut with 4-0 surgical suture. The mice were monitored for at least 48 hours post-surgery. The AAVs included either U6-sgRNA-hSyn-Cre-2A-EGFP-KASH AAV with either a lacZ (control) gRNA or a gRNA directed against the *Bdnf* promoter I (P1g3) or intronic region (IEg3). *Incubation of cocaine craving, day 21*:After 20 days of abstinence in the home cage, mice were tested for incubation of craving on day 21. *Food self-administration with Bdnf knockdown*: To ensure that the experimental *Bdnf* knockdown did not alter the ability to press a lever in the incubation test, a subset of the mice were then examined for food self-administration under the previously established FR5TO20 schedule of reinforcement across three baseline 1 hr testing sessions. To then assess cognitive flexibility, subjects underwent the lever reversal procedure, in which the previously active lever was reassigned to be inactive (noncontingent) lever, and the inactive lever was reassigned to be active lever (FR5TO20 schedule with food reward). Mice were tested for an additional four 1 hr sessions with the reversed lever conditions. Subjects were sacrificed after behavioral testing, and brain sections were processed for injection validation.

### Dopaminergic nuclei isolation

We used a variation of the published INTACT protocol (Mo et al., 2015). All the following steps were performed on ice or in the cold room. The VTA samples were defrosted and dounce-homogenized using first a loose pestle, then a tight pestle in 1.5ml of homogenization buffer (0.25M sucrose, 25mM KCl, 5mM MgCl2, 20mM Tricine-KOH) supplemented with 1mM DTT, 0.15mM spermine, 0.5mM spermidine, 172g/L kynurenic acid, 5mM sodium butyrate and EDTA-free protease inhibitor. A 5% IGEPAL-630 solution was added to bring the homogenate to 0.3% IGEPAL-630, and the samples were further dounced. These samples were then passed through a 40um strainer and mixed with 1.5ml Working Solution (1:5 ratio of 150mM KCl, 30mM MgCl2, 120mM KOH, pH 7.8 Diluent and Optiprep Density Gradient Medium). These were then underlaid with a gradient of 30% and 40% iodixanol and centrifuged at 10,000 rpm for 19 minutes in a Sw41Ti motor swinging bucket centrifuge at 4⁰C. The separated nuclei at the interface of the gradients were collected and further processed for dopaminergic nuclear enrichment.

For bulk RNA-seq and for HiC on Accessible Regulatory DNA (HiCAR) chromatin conformation experiments, the nuclear fraction was precleared by incubation with 15µl of protein G magnetic Dynabeads (Thermo Fisher Scientific Cat#10004D) for 15 min. After removing the Dynabeads using a magnet, the samples were diluted in homogenization buffer and incubated with 10µl of 0.2mg/ml rabbit anti-GFP antibody (Thermo Fisher Scientific Cat# G10362, RRID:AB_2536526) for 1 hour, end-to-end rotated at 4⁰C. 60µl of the Dynabeads were then added to these samples and they were end-to-end rotated at 4⁰C for 1 hour. To improve the yield of enrichment, the bead-nuclei mixture was place on the magnet for 30sec and then completely resuspended by inversion. This was repeated 5-7 times, after which they were placed on the magnet for 5 min. 1mL of the supernatant was collected as the Unbound Fraction (UF) and centrifuged at 2000 RPM for 5 min to pellet the nuclei. The beads were then washed 3 times with homogenization buffer, followed by one wash in 6mL of homogenization buffer. The 6mL were then sequentially applied to the magnet until all beads had been isolated. This was the immunoprecipitated fraction (IP).

For single nucleus RNA sequencing (snRNA-seq) experiments, the nuclei fraction was pelleted and resuspended in MACS nuclei separation buffer (Miltenyi Biotec, 130-128-024). The samples were then incubated with 50µl of Miltenyi MACS anti-GFP Microbeads (Miltenyi Biotec, 130-091-125) and end-to-end rotated at 4⁰C for 15 minutes. The GFP+ nuclei were then enriched by following the Miltenyi MACS magnetic nuclei separation protocol using MS columns (Miltenyi Biotec, 130-042-201). Nuclei per sample were counted using the Countess 3 FL Automated Cell Counter (Invitrogen, AMQAF2000) with Hoechst staining (Thermo Fisher Scientific Cat# H1399, Dilution 1:1000). 75-100k nuclei per animal were then pelleted, resuspended in 1X PBS, and then incubated with 5µl of oligos from the ScalePlex Oligo Plate (Scale Biosciences 1064662) to tag and label nuclei from individual animals. These nuclei were then fixed using the ScaleBio Sample Fixation Kit (Scale Biosciences 202003), pooled together, and went through a series of wash steps using the Wash Buffer (Scale Biosciences 202100001). The nuclei were then store at-80°C until all samples were collected and fixed. To eliminate any batch effects that may arise due to nuclear collection and fixation at different times, each pool consisted of nuclei from one animal of each treatment condition.

### Bulk RNAseq library preparation and sequencing

RNA for bulk RNA-seq was obtained using the RNAqueous – Micro Total RNA Isolation kit (Invitrogen, AM1931). RNA purity was measured to ensure that samples had A260/280 and A260/230 > 1.9. RNA was then polyA enriched and 150bp paired-end sequencing was performed by Novogene, Inc on a NovaSeq X Plus machine.

### Bulk RNAseq data processing

Quality scoring of reads, adapter trimming, read mapping to the mm10 reference genome, generation of bigwig files and read counts files were done using the nf-core rnaseq pipeline: nf-co.re/rnaseq/3.22.2.Differential gene expression was calculated using DESeq2.

### Single nucleus RNAseq library preparation and sequencing

The snRNAseq library was prepared from a total of 20 samples (isolated and fixed dopaminergic nuclei from 4 Saline AD1 animals, 6 Saline AD14 animals, 4 Cocaine AD1 animals, and 6 Cocaine AD14 animals, between the ages of P60-P90). Equal numbers of male and female mice were included in each experimental condition. On the day of library preparation, the pooled nuclei were thawed on ice and each pool was counted using the Countess 3 FL Automated Cell Counter (Invitrogen, AMQAF2000). 75,000-100,000 nuclei were used per animal.

For library preparation, the ScaleBio Single Cell RNA Sequencing Kit v1.1 (Scale Biosciences 950884) was used according to manufacturer’s instructions. Nuclei from each pool of fixed and tagged nuclei were loaded at 6,500 nuclei per well to the 96-well RT Barcode plate to add the RT barcode and UMI onto the transcripts during reverse transcription for cDNA synthesis. The nuclei from these wells were then pooled by centrifugation using the ScaleBio Sample Collection Funnel, following which they were redistributed across the 384-well Ligation Barcode Plate for the addition of ligation adaptors and barcodes to the UMI-RT barcoded transcript. This ligation adaptor contains the TruSeq Read 1 sequence. The nuclei were pooled again using the ScaleBio Sample Collection Funnel. A total of 1600 nuclei were distributed per well of the 96-well Final Distribution Plate. In each well, second strand synthesis was performed, followed by a cleanup step to break down the nuclei. The transcripts were then tagmented to the Nextera Read 2 sequence, after which an indexed PCR was performed to amplify the library, as well as to add 96 unique Index i5 barcodes and 4 unique i7 indices. An aliquot of this unpurified RNA library was further PCR-amplified to enrich the ScalePlex constructs and create the ScalePlex library, which can be differentiated from the RNA library by a unique i7 Index barcode. Finally, the libraries were cleaned up to select for 300-500bp fragments using 0.8X SPRIselect beads (Beckman Coulter B23317). 6µl from each of the 96 RNA libraries were pooled and cleaned using the SPRIselect beads, while a total of 40µl of the ScalePlex library was cleaned up using the SPRIselect beads. The average fragment size of the final libraries was quantified on a Agilent TapeStation 4200. The RNA and ScalePlex libraries were sequenced by the Duke Sequencing Core on a 25B flowcell at a 10:1::RNA:ScalePlex library ratio and sequenced on Novaseq X Plus (Illumina) to a target depth of 10,000 reads per nucleus.

### snRNAseq data processing

Base calls were converted to fastq files and demultiplexed by Index1 barcode by the Duke Sequencing Core. Combinatorial barcode demultiplexing, barcode processing, adapter trimming, read mapping to the mm39 reference genome, single-nuclei counting, and generation of the feature-barcode matrices were done using the Nextflow ScaleRNA pipeline v1.6.3 (https://github.com/ScaleBio/ScaleRna). The filtered count matrices were brought into Seurat for downstream analysis.

For quality control, a UMI-gene cutoff of 200-5000 UMIs and 200-20,000 genes for each sample was performed. Barcodes with a mitochondrial reads percentage greater than 5 were removed. After quality control filtering, the dataset contained 46,612 nuclei across all samples, with a median of 5665 UMIs and 2652 genes per nucleus. The count matrices were log2 normalized, centered, and scaled using a scaling factor of 10,000. The top 2000 most variable genes were identified using dispersion and mean expression thresholds. Principal component analysis (PCA) was performed, followed by an elbow plot to determine the true dimensionality of the data, which for this dataset was PC 20-21. Dimensionality reduction by UMAP was then performed for 30 dimensions, followed by a clustering approach using the Louvain algorithm. Once the dopamine cluster was identified, it was subset into a new Seurat object and the pre-processing steps of normalization, centering, scaling, PCA, and dimensionality reduction using UMAP were repeated. The gene expression profiles of the dopamine subclusters were compared to those of dopamine subpopulations identified in other studies (Poulin et al., 2020) to identify our subclusters.

### Differential gene expression analysis

To identify genes differentially expressed in the cocaine AD14 group compared to cocaine AD1 group in the dopamine cells, we used MAST (Finak et al., 2015) to perform zero-inflated regression analysis by fitting a linear mixed model (LMM). We considered as significant genes that had a log-transformed fold change of expression of at least 0.25 (20% difference) and FDR<0.05.

### HiCAR method

For the AD14 saline and cocaine DAT HiCAR libraries, 50,000 nuclei/sample was used. We had 4 biological replicates per treatment condition (2 male, 2 female) and used DA neurons isolated from 1 mouse per library. HiCAR was performed as previously described (Wei et al., 2022). Briefly, Nuclei and Tn5 transposase were incubated with agitation at 37°C for 1 hour. Tagmented DNA was digested with MseI restriction enzyme (NEB, cat #R0525S) alongside splint2 oligonucleotide (**Table S2**). In-situ ligation was performed using T4 DNA ligase (NEB, cat #M0202L) at room temperature for 4 hours. Following ligation, samples were reverse-crosslinked overnight at 68°C. DNA was ethanol precipitated at-80°C for 30-60 min. DNA was then digested with NIaIII restriction enzyme (NEB, #R0125S) at 37°C for 1 hour and cleaned up using 0.9X volume of SPRI beads (Beckman, cat #B23319). DNA concentration was measured using a Qubit instrument and adjusted to a concentration of 1 ng/µL. A second round of ligation was performed using T4 DNA ligase incubated at RT for 2 hours. DNA was purified using the Zymo DNA clean and concentrator kit (Zymo, cat #D4029) and eluted using 10mM Tris-HCl, pH 8.0. A PCR reaction mixture was set up, using unique Nextera-pcr-i7-primers and NEB i5 primers to amplify and index samples (**Table S3**). Resulting libraries were checked for fragment size using a bioanalyzer, after which they were pooled and sequenced. Sequencing was performed by BGI Genomics on the DNBSeq sequencing platform.

### HiCAR data processing

HiCAR fastq files were processed using the Nextflow nf-core (Ewels et al., 2020) HiCAR pipeline (Wei et al., 2022) using Nextflow v24.10.5 Jianhong/HiCAR ver2rc (OU et al., 2022). The container engine used to run the Jianhong/HiCAR pipeline was singularity, and default pipeline parameters were used. QC metrics are shown in **Table S4.**

### Homer

Using bedtools, the TAD boundaries bed file and the ATAC-seq bed file generated by the nf-core HiCAR pipeline from the saline DAT cells were intersected to generate a new bed file containing the ATAC peaks within TADs of the differentially expressed genes discovered from the snRNA-seq data. This new bed file was used as the input for HOMER, a PWM-based method for motif enrichment, to identify TF motifs enriched within these ATAC peaks, with the background file being all ATAC peaks in saline DAT cells.

## Statistical analyses

For experiments other than sequencing, data were analyzed using standard t-test, nested t-test, one-way ANOVA, or two-way ANOVA, as appropriate to the data structure and as described in their respective results section. All analyses were performed on GraphPad Prism 10 (RRID:SCR_002798). Depending on data structure either P value or FDR <0.05 was considered significant.

## Data availability

All RNAseq, scRNAseq, and HiCAR data will be deposited in a publicly available database at the time of manuscript acceptance.

## Code availability

Scripts used for this analysis will be deposited in a publicly available database at the time of manuscript acceptance.

### Egr1 Immunocytochemistry

Flash-frozen and cryosectioned brain slices were fixed in 4% formaldehyde, following which they were blocked and permeabilized with blocking solution (10% Normal Goat serum, 0.3% Triton X-1000, PBS). They were then incubated with primary antibodies (rabbit anti-Egr1 antibody: Cell Signaling Cat#4153, Dilution 1:1000; chicken anti-TH antibody: Abcam Cat#ab76442, Dilution 1:1000) suspended in blocking solution overnight at 4°C, following which they were washed with PBS and incubated with secondary antibodies (goat anti-rabbit 594: Invitrogen Cat#A11012, Dilution 1:500; goat anti-chicken 488: Invitrogen Cat#A11039, Dilution 1:500) in blocking solution for an hour at room temperature. Sections were washed with PBS thrice and nuclei were then labelled with Hoechst (1:1000 Thermo Fisher Scientific, H1399), and coverslipped and mounted in Vectashield (VectorLabs H1000-10).

## Results

### Bdnf is expressed in multiple cell types within the VTA

Prior single cell sequencing studies of microdissected VTA have shown that *Bdnf* mRNA is expressed in glutamatergic as well as dopaminergic neurons within this structure (Phillips et al., 2022). To establish an assay for quantitatively measuring *Bdnf* mRNA in VTA with cell-type precision, we performed multiplex RNAScope single-molecule fluorescent *in situ* hybridization (FISH) on coronal sections of adult mouse VTA. We identified dopaminergic neurons using a probe targeting the RNA encoding the dopamine transporter, DAT (*Slc6a3*), and glutamatergic neurons using a probe targeting the RNA encoding the vesicular glutamate transporter, VGLUT2 (*Slc17a6*). A nuclear dye was used to identify all cells in the section. We used a *Bdnf* probe that targets the coding exon of *Bdnf,* allowing us to see all *Bdnf* transcripts regardless of which of the multiple *Bdnf* promoters initiates transcription (West et al., 2014). We imaged and separately analyzed cells from the medial and lateral VTA (**Fig. 1A,B**), because prior studies have indicated distinct roles for these regions and their interconnected circuits in reward (Chaudhury et al., 2013; Lammel et al., 2011; Morales & Margolis, 2017).

**Figure 1:**
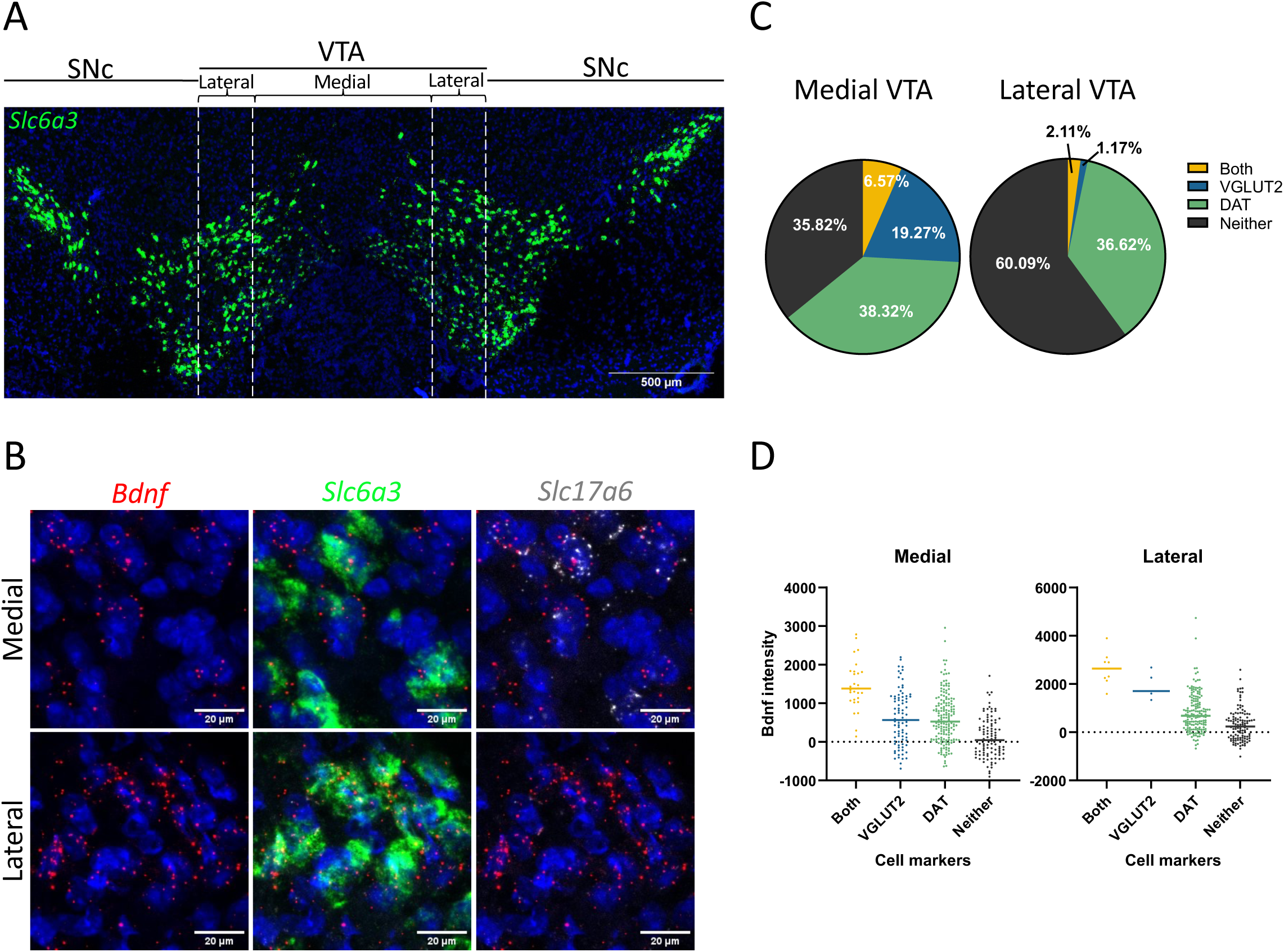
***Bdnf* is expressed in multiple cell types within the VTA. A)** Representative RNA in situ hybridization image of the VTA, with the dotted white lines defining the boundaries between medial and lateral VTA. (VTA: Ventral Tegmental Area, SNc: Substantia nigra pars compacta) **B)** Quantification of cells positive for the *Slc17a6* probe (VGLUT2), the *Slc6a3* probe (DAT), both probes (Both), or neither probe (Neither), in the medial (left) and the lateral (right) VTA. **C)** Representative RNA in situ hybridization images from the medial (top panels) and the lateral (bottom panels) VTA showing colocalization of *Bdnf* signal (red) with Slc6a3 (green) and/or Slc17a6 (white). **D)** Quantification of Bdnf signal in different cell types. Statistics: n=3 mice, 50-70 cells sampled per animal, One-way ANOVA with post-hoc being Tukey’s multiple comparison; ns = not significant, *p<0.05, **p<0.01, ***p<0.001, ****p<0.0001. Error bars indicate standard deviation (SD).

In the medial VTA, 38.32 ± 3.17% of all cells were DAT neurons (positive for *Slc6a3*) and 19.27 ± 2.61% were VGLUT2 neurons (positive for *Slc17a6*) (**Fig 1B,C**; mean ± SEM, n = 3 mice). We observed *Bdnf* expression in both cell populations, and we saw similar levels of *Bdnf* per cell in both cell types in the medial VTA (**Fig 1D**). In the lateral VTA, DAT cells comprised 36.62 ± 10.16% of all cells whereas, consistent with prior reports (Morales & Margolis, 2017), there were very few VGLUT2 cells (1.17 ± 0.85%) (**Fig 1B,C**). Thus, both DAT and VGLUT2 cells expressed *Bdnf* in medial and lateral VTA with different relative proportions in each subregion.

In both medial and lateral VTA we also observed a small percentage of cells that were positive for both *Slc6a3* and *Slc17a6.* Combinatorial cells that release multiple neurotransmitters including both dopamine and glutamate have been identified in many subcortical regions including the VTA (Morales & Margolis, 2017). These dual DAT/VGLUT2 neurons comprised only 6.57% of all cells in the medial VTA and 2.11% in lateral VTA (**Fig 1C**), however they showed the highest overall levels of *Bdnf* (**Fig 1D**). Finally, we did observe low, but detectable, *Bdnf* expression in some cells that expressed neither *Slc6a3* nor *Slc17a6* (**Fig 1D**). These may be non-neuronal cells, such as astrocytes and microglia, which are known to express low levels of *Bdnf* (Zhang et al., 2014). The failure to detect *Bdnf* in non-neuronal cells in single cell sequencing studies of the VTA (Phillips et al., 2022) may be a consequence of the very low expression of *Bdnf* in these cells and the limited sensitivity of scRNAseq (Qiu, 2020). Due to the low number of the dual *Slc6a3/Slc17a6* cells and the unknown identity of the cells that express neither marker, these populations were not considered further in this study.

### Time-dependent upregulation of Bdnf in VTA DAT neurons upon cocaine abstinence

To determine whether abstinence after chronic cocaine drives upregulation of *Bdnf* mRNA in specific cell types within the VTA, we quantified VTA *Bdnf* expression by FISH after a period of repeated cocaine exposure. Adult male and female mice underwent repeated saline or cocaine (20mg/kg i.p.) injections daily for 7 days. We then harvested their brains either 1 day (abstinence day 1, AD1) or 14 days (AD14) after the final saline or cocaine injection to process them for multiplex FISH as above **(Fig. 2)**. To permit 2-way statistical comparisons across time and treatment we averaged the *Bdnf* expression across all cells within each given cell population (medial or lateral, DAT or VGLUT2) in each single mouse.

**Figure 2:**
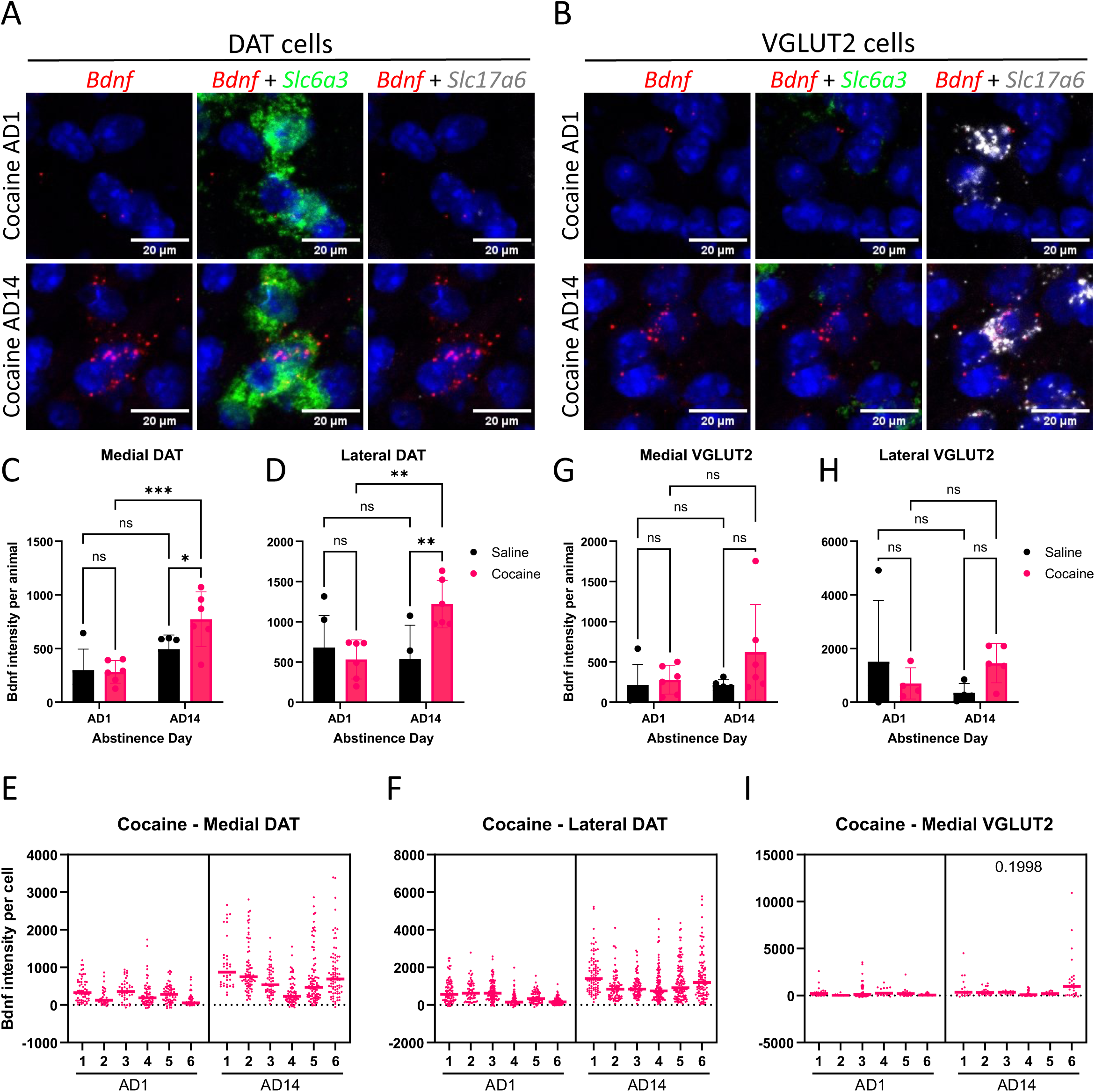
***Bdnf* is upregulated in VTA DAT neurons upon forced cocaine abstinence.** A) Representative RNA *in situ* hybridization images of *Bdnf* signal in medial DAT cells, indicated by the presence of *Slc6a3* signal and absence of *Slc17a6* signal, at Cocaine AD1 and AD14.B) Representative RNA *in situ* hybridization images of *Bdnf* signal in medial VGLUT2 cells, indicated by the absence of *Slc6a3* signal and presence of *Slc17a6* signal, at Cocaine AD1 and AD14. C) Average *Bdnf* signal in the medial DAT cells in each animal upon cocaine FA. D) Average *Bdnf* signal in the lateral DAT cells in each animal upon cocaine FA. E) Average *Bdnf* signal in the medial VGLUT2 cells in each animal upon cocaine FA. F) Average *Bdnf* signal in the lateral VGLUT2 cells in each animal upon cocaine FA. G*) Bdnf* signal in medial DAT cells of each animal in Cocaine AD1 and AD14 groups. H*) Bdnf* signal in lateral DAT cells of each animal in Cocaine AD1 and AD14 groups. I) *Bdnf* signal in medial DAT cells of each animal in Cocaine AD1 and AD14 groups. Statistics: n=4-7mice, 50-70 cells sampled per animal, 2C-D: Two-way ANOVA with post-hoc Tukey’s multiple test, 2E-G: Nested t-test; ns = not significant *p<0.05, **p<0.01, ***p<0.001, ****p<0.0001. Error bars indicate standard deviation (SD).

At AD1, the levels of *Bdnf* were not significantly different between mice that received cocaine versus saline whether we compared the medial or lateral DAT or VGLUT2 neurons **(Fig. 2A-F)**. These data show that repeated cocaine exposure alone does not increase *Bdnf* expression in VTA neurons. We then asked if *Bdnf* expression is induced over the AD14 period of abstinence after cocaine. Here we compared *Bdnf* expression in DAT or VGLUT2 neurons after 1d of abstinence after cocaine or saline to the levels observed after 14d of abstinence **(Fig. 2A-F**). These data showed a significant increase in *Bdnf* expression in DAT neurons in both the medial (**Fig 2C**) and the lateral (**Fig 2D**) VTA after 14d of abstinence from cocaine compared either with the levels seen after 1d of cocaine abstinence or with the levels seen 14d after saline injections (**Fig 2C**:Two-way ANOVA, Time F_(1, 18)_ = 19.07, p = 0.0004; Treatment: F_(1, 18)_ = 2.796, p = 0.1118; Interaction: F_(1, 18)_ = 3.530, p = 0.0766; **Fig 2D**: Two-way ANOVA, Time: F_(1, 18)_ = 3.452, p = 0.0796; Treatment: F_(1, 18)_ = 3.351, p = 0.0838; Interaction: F_(1, 18)_ = 8.109, p = 0.0107). The significant induction of *Bdnf* at the single cell level in medial and lateral DAT neurons was even more apparently when we graphed the distribution of single neuron *Bdnf* levels in each of the mice that experience 1d versus 14d of cocaine abstinence (**Fig 2E**: Nested t-test, t=4.170, df=10, p = 0.0019; **Fig 2F**: Nested t-test, t=4.448, df=10, p = 0.0012).Thus *Bdnf* induction in VTA DAT neurons is dependent both on a past history of cocaine exposure and on a prolonged period of abstinence. By contrast we saw no induction of *Bdnf* after cocaine abstinence in either the medial (**Fig 2G**) or lateral (**Fig 2H**) VGLUT2 cells when we either considered the average expression per animal or the per cell *Bdnf* levels in individual mice **(Fig. 2I)**. (**Fig 2G**: Two-way ANOVA, Time: F_(1, 18)_ = 1.303, p = 0.2686; Treatment: F_(1, 18)_ = 2.487, p = 0.1322; Interaction: F_(1, 18)_ = 1.288, p = 0.2713; **Fig 2H**: Two-way ANOVA, Time: F_(1, 13)_ = 0.1160, p = 0.7389; Treatment: F_(1,13)_ = 0.06298, p = 0.8058; Interaction: F_(1, 13)_ = 2.625, p = 0.1292; **Fig. 2I**: Nested t-test. t=1.373, df=10, p = 0.1998)

### Promoter I and an intragenic enhancer mediate the induction of Bdnf mRNA in DAT neurons upon cocaine abstinence

Levels of *Bdnf* are tightly controlled at the transcriptional level by the activation of transcription factors bound to at least eight promoters as well as an intragenic enhancer (IE) within the *Bdnf* gene body (**Fig. S1A)** (Kim et al., 2010; Tuvikene et al., 2021; West et al., 2014). No matter which promoter is used, all of the *Bdnf* transcripts encode the same mature peptide (West et al., 2014). Of the transcript variants, *BdnfI* (initiating at promoter I, P1) and *BdnfIV* (initiating at promoter IV, P4) are the most highly expressed in the brain (West et al., 2014). Each promoter and enhancer is bound by different complements of transcription factors, thus this complex regulatory architecture allows cell-type and stimulus-specific control over *Bdnf* transcription (Griffith et al., 2024). Prior studies using quantitative PCR from tissue punches from rats have indicated that *BdnfI* is the most highly expressed *Bdnf* transcript in VTA and the one most strongly induced by cocaine abstinence (Schmidt et al., 2012). We reasoned that if we could use our knowledge of the transcriptional regulation of *Bdnf* to selectively block its induction in the VTA during cocaine abstinence, then we could determine the functional consequence of this gene’s regulation for drug seeking behaviors.

To control transcription of *Bdnf* via specific regulatory elements, we designed guide RNAs (gRNAs) to recruit the CRISPR interference (CRISPRi) repressor dCas9-KRAB to *Bdnf* P1, the IE, or P4. We selected guides by validating their efficiency to repress specific *Bdnf* transcripts in mouse embryonic cortical neuron cultures. Because embryonic neurons express low levels of *Bdnf* relative to the adult brain (Maisonpierre et al., 1990), we used KCl membrane depolarization as a stimulus to robustly induce *Bdnf* transcription (Tao et al., 1998). As a control for off-target effects of dCas9-KRAB expression, for comparison we used a non-targeting gRNA with a sequence complementary to the bacterial gene *lacZ*.

When P1 gRNAs were co-expressed with dCas9-KRAB (**Fig S1B**), they significantly suppressed membrane depolarization-induced expression of *BdnfI* relative to cells expressing the lacZ gRNA (*BdnfI*: Two-way ANOVA, gRNA: F_(2, 30)_ = 29.15, p<0.0001; Stimulation: F_(1, 30)_ = 137.1, p<0.0001; Interaction: F_(2, 30)_ = 26.88, p<0.0001). *BdnfIV* expression was unaffected showing the specificity for the *BdnfI* promoter (*BdnfIV*: Two-way ANOVA, gRNA: F_(2, 30)_ = 0.6486, p = 0.5299; Stimulation: F_(1, 30)_ = 83.01, p<0.0001; Interaction: F_(2, 30)_ = 0.5850, p = 0.5633). Conversely, co-expression of *Bdnf* P4 targeted gRNAs with dCas9-KRAB (**Fig S1C**) significantly suppressed induction of *BdnfIV* while having no effect on *BdnfI* induction (*BdnfI*: Two-way ANOVA, gRNA: F(2, 18) = 0.1975, p = 0.8225; Stimulation: F(1, 18) = 188.0, p<0.0001; Interaction: F(2, 18) = 0.2250, p = 0.8007; *BdnfIV*: Two-way ANOVA, gRNA: F_(2, 17)_ = 8.799, p = 0.0024; Stimulation: F_(1, 17)_ = 250.7, p<0.0001; Interaction: F_(2, 17)_ = 8.543, p = 0.0027). When IE gRNAs were co-expressed with dCas9-KRAB (**Fig S1D**), we observed significantly reduced expression of both *BdnfI* and *BdnfIV* raising the possibility that this enhancer may co-operate with both the upstream (I-III) and downstream (IV-VII) *Bdnf* promoter clusters (*BdnfI*: Two-way ANOVA, gRNA: F_(2, 29)_ = 5.447, p = 0.0098; Stimulation: F_(1, 29)_ = 234.1, p<0.0001; Interaction: F_(2, 29)_ = 5.406, p = 0.0101; *BdnfIV*: Two-way ANOVA, gRNA: F_(2,29)_ = 5.514, p = 0.0093; Stimulation: F_(1, 29)_ = 389.9, p<0.0001; Interaction: F_(2, 29)_ = 5.350, p = 0.0105). Importantly, expression of any of these *Bdnf* targeted gRNAs had no effect on KCl-dependent induction of the activity-regulated genes *Pcsk1* or *Nr4a2* (Tyssowski et al., 2018) (**Fig. S1B-D),** demonstrating that the inhibition of *Bdnf* by CRISPRi was specific and not secondary to a general failure of depolarization-induced signaling cascades (**Fig S1B** *Pcsk1*: Two-way ANOVA, gRNA: F_(2, 30)_ = 0.7636, p = 0.4748; Stimulation: F_(1, 30)_ = 132.5, p<0.0001; Interaction: F_(2, 30)_ = 0.6599, p = 0.5242; **Fig S1C** *Pcsk1*: Two-way ANOVA, gRNA: F_(2, 18)_ = 0.5735, p = 0.5735; Stimulation: F_(1, 18)_ = 84.24, p<0.0001; Interaction: F_(2, 18)_ = 0.5774, p = 0.5714; **Fig S1D** *Nr4a2*: Two-way ANOVA, gRNA: F_(2, 30)_ = 3.360, p = 0.0482; Stimulation: F_(1,30)_ = 1187, p<0.0001; Interaction: F_(2, 30)_ = 3.433, p = 0.0454).

To determine the contributions of these gene regulatory elements to *Bdnf* upregulation after chronic cocaine in the VTA, we stereotaxically injected the VTA of conditional dCas9-KRAB mice (Gemberling et al., 2021) with AAVs expressing Cre recombinase and GFP under control of the neuronal *Syn1* promoter as well as one of the gRNAs. We then gave mice repeated saline or cocaine injections as described above. Brains were harvested on AD14 and processed for FISH. The cocaine-dependent upregulation of *Bdnf* at AD14 was observed in mice with the control *lacZ* gRNA virus, confirming that this regulation is not impaired by the stereotaxic surgery, AAV infection, or the expression of Cre, GFP, and dCas9-KRAB (**Fig S2A**: Unpaired t-test, t=16.51, df=444; **Fig S2B**: Unpaired t-test, t=4.899, df=590).

In both the medial and the lateral DAT cells, recruiting dCas9-KRAB to either *Bdnf* P1or the IE significantly reduced *Bdnf* relative to mice infected with the control nontargeting *lacZ* gRNA (**Fig 3B-E**) (**Fig 3B**: One-way ANOVA, F_(3,10)_ = 1.027, p = 0.4217; **Fig 3C**: Nested one-way ANOVA, F_(3,10)_ = 8.285, p = 0.0046; **Fig 3D**: One-way ANOVA, F_(3,12)_ = 0.7128, p = 0.5629; **Fig 3E**: Nested one-way ANOVA, F_(3,12)_ = 10.65, p = 0.0011). This was observed when the data were quantified as either average *Bdnf* signal per animal per condition (**Fig 3B, D)** or as *Bdnf* signal per cell per animal in each condition (**Fig 3C, E)**. When *Bdnf* P4 was targeted by dCas9-KRAB, there was no significant effect (**Fig 3 B-E**). *Bdnf* was also induced in medial VTA VGLUT2 neurons following 14d of cocaine abstinence, however in these cells all three gRNAs (P1, IE, P4) significantly blocked *Bdnf* induction (**Fig. S3)**. From these experiments we chose to use P1 and IE gRNAs to block *Bdnf* induction during abstinence after repeated cocaine.

**Figure 3:**
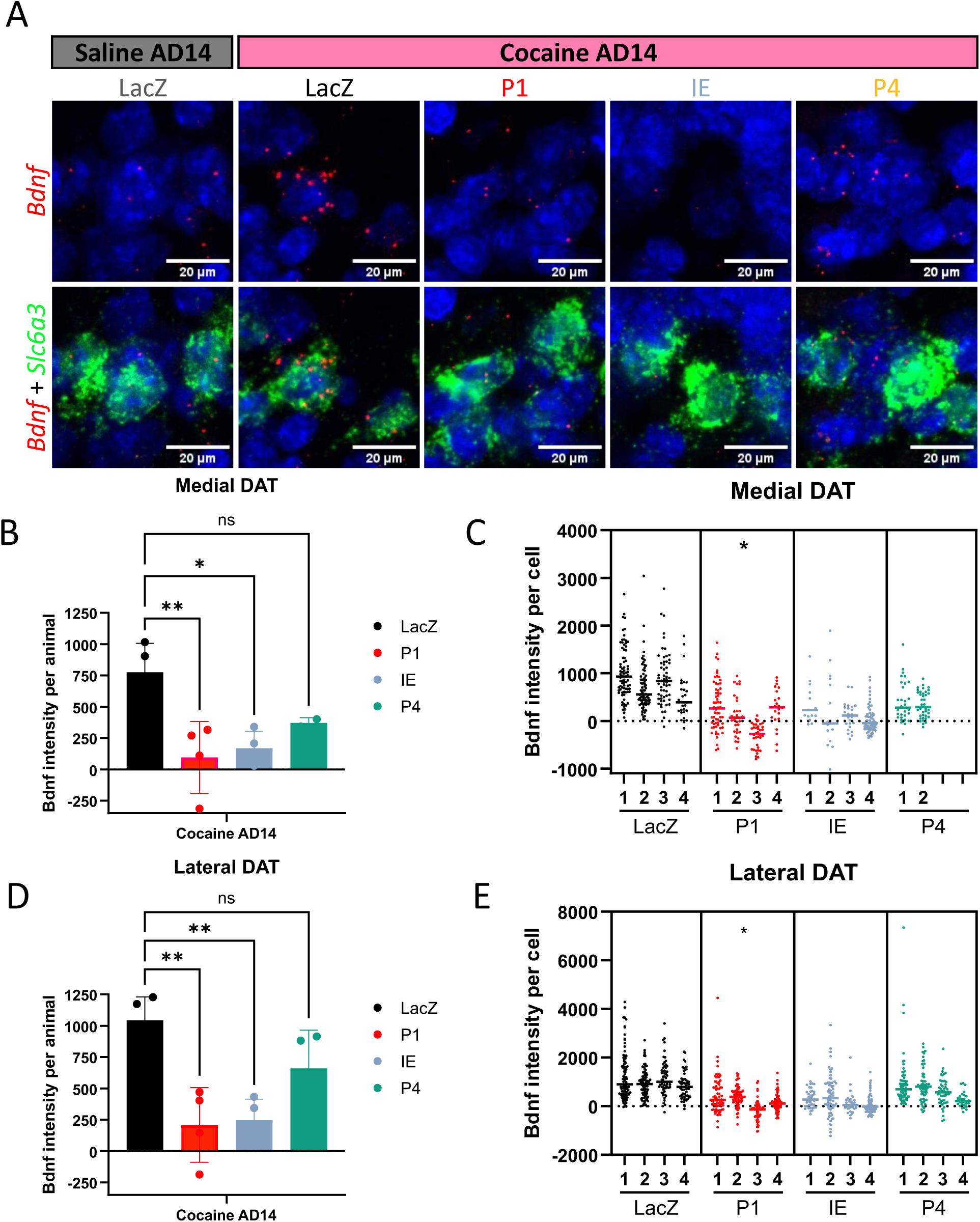
dCas9-KRAB-mediated repression of *Bdnf* promoter I and the intragenic enhancer reduces Bdnf upregulation upon forced cocaine abstinence. Representative RNA *in situ* hybridization images of *Bdnf* signal (red) in medial DAT cells (*Slc6a3* in green) from the VTA of mice injected with LacZ, P1, IE, or P4 gRNA AAV. A) Quantification of *Bdnf* signal in medial DAT cells as average *Bdnf* signal per mouse injected with LacZ, P1, IE, or P4 gRNA AAV, treated with cocaine and harvested at AD14. B) Quantification of *Bdnf* signal in medial DAT cells as *Bdnf* signal per cell per mouse injected with LacZ, P1, IE, or P4 gRNA AAV, treated with cocaine and harvested at AD14. C) Quantification of *Bdnf* signal in lateral DAT cells as average *Bdnf* signal per mouse injected with LacZ, P1, IE, or P4 gRNA AAV, treated with cocaine and harvested at AD14. D) Quantification of *Bdnf* signal in lateral DAT cells as *Bdnf* signal per cell per mouse injected with LacZ, P1, IE, or P4 gRNA AAV, treated with cocaine and harvested at AD14. Statistics: n=4 mice, 50-70 cells sampled per animal, 2B,D: One-way ANOVA with post-hoc test being Tukey’s multiple test, 2C,E: Nested One-way ANOVA with post-hoc test being Tukey’s multiple test; ns = not significant *p<0.05, **p<0.01, ***p<0.001, ****p<0.0001. Error bars indicate standard deviation (SD).

### Repression of Bdnf after cocaine self-administration blocks incubation of cocaine craving

Prior studies showed that artificially increasing BDNF protein in the VTA promotes cocaine seeking (Lu et al., 2004); however, it was not known whether *Bdnf* transcription in the VTA during abstinence is required for the incubation of cocaine seeking. To test this, we used the incubation of cocaine seeking paradigm (Grimm et al., 2001), in which mice are first trained to self-administer intravenous cocaine and then tested for their drug seeking behavior over a period of forced abstinence (**Fig S4A**). Conditional dCas9-KRAB mice were provided access to self-administer cocaine (0.3 mg/kg/infusion) in operant chambers under a fixed ratio 5, time out 20 sec (FR5TO20) schedule of reinforcement. After 10 days of self-administration, on day 1 of abstinence (AD1), the mice were placed in the operant chamber where they were allowed to lever press though no cocaine was available. We quantified active lever press responses paired with the cue light according to the FR5TO20 schedule but in the absence of cocaine infusions, as well as inactive lever responses. Immediately following this test, mice were stereotaxically injected in the VTA with AAVs expressing Cre along with one of the gRNAs tested above. They received either a *lacZ* gRNA as control or gRNAs targeting *Bdnf* P1 **(Fig 4)** or the intragenic enhancer **(Fig 5)** to induce *Bdnf* knockdown (**Fig S4B,C**). Thereafter, they underwent the abstinence period in their home cage across 19 days to allow for full expression of the virus. On day 21 of abstinence (AD21), the mice were placed back in the operant chamber and again we quantified active and inactive lever press responses in the absence of cocaine infusions.

**Figure 4:**
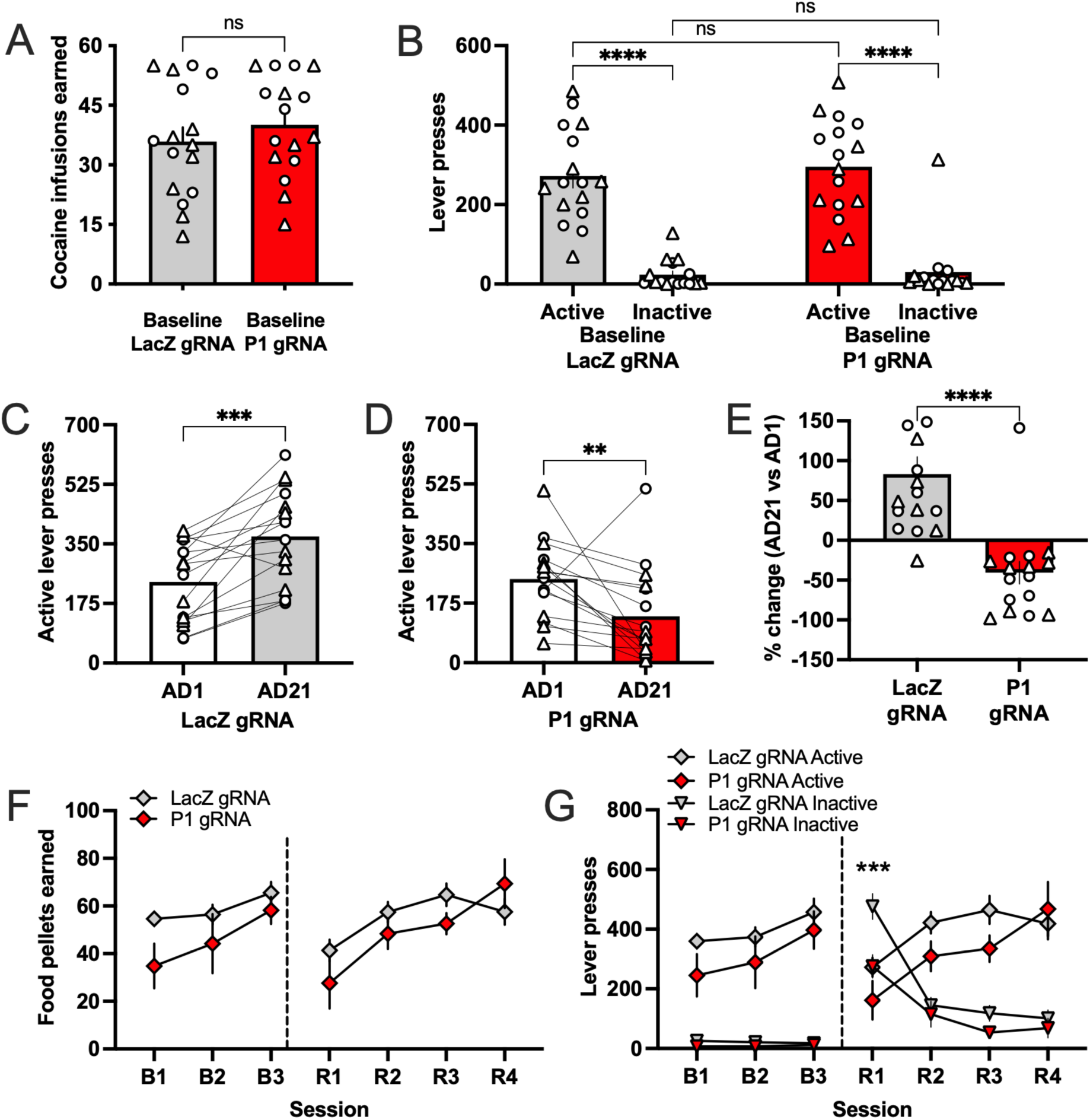
dCas9-KRAB-mediated repression of *Bdnf* promoter I reduces cocaine seeking behaviors in mice. A) Number of cocaine infusions earned prior to injection of either the control LacZ gRNA AAV or the knockdown P1 gRNA AAV (n=16/group; 8 males/group denoted with circles, 8 females/group denoted with triangles). B) Active and Inactive lever presses for mice prior to injection of the control LacZ gRNA or knockdown P1 gRNA AAV. Both groups demonstrated preference for the active lever, and there were no differences in the active or inactive lever responding between groups. C) Mice injected into the VTA with the control LacZ gRNA exhibited a robust incubation of craving response, with greater active lever presses on day 21 (AD21) compared to day 1 (D1) of forced abstinence. (n=16; 8 males, 8 females). D) Mice injected into the VTA with the knockdown P1 gRNA did not exhibit incubation of cocaine craving. Rather, they displayed a reduction in active lever pressing on day 21, compared to day 1. (n=16; 8 males, 8 females). E) In comparing the percent change in active lever presses from day 21 to day 1, the control LacZ and P1 gRNA groups exhibited opposing responses (n=16/group; 8 males/group, 8 females/group). F) No differences were found in the number of food pellets earned by control LacZ gRNA and knockdown P1 gRNA mice during baseline (B1-3) and reversal sessions (R1-4) of the food lever reversal task (n=5-11/group; control, 5 males, 6 females; knockdown P1, 3 males, 2 females). G) Number of active and inactive lever presses in control and knockdown mice during the food lever reversal task. Whereas the groups did not differ in responses on the food-associated active lever, the inactive lever responding was greater on day 1 of reversal for the control LacZ group, as compared to the knockdown P1 gRNA group. Statistics: ns = not significant, **p<0.01, ***p<0.001, ****p<0.0001. Error bars indicate standard error mean (SEM).

**Figure 5:**
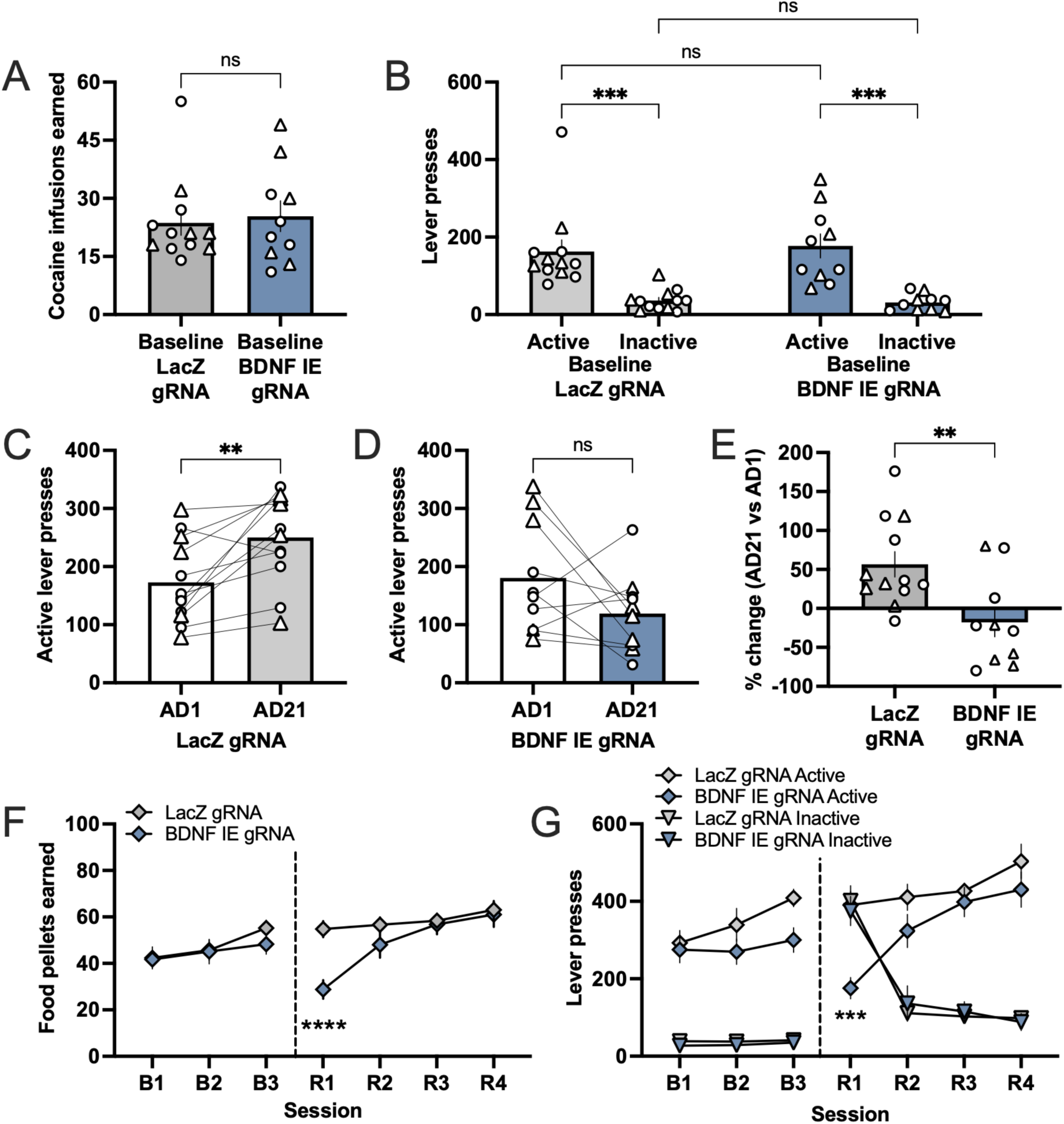
dCas9-KRAB-mediated repression of *Bdnf* intragenic enhancer (IE) reduces cocaine seeking behaviors in mice. A) Number of cocaine infusions earned prior to injection of either the control LacZ gRNA AAV or the knockdown IE gRNA AAV (n=5-7/group; control, 7 males denoted with circles, 5 females denoted with triangles; knockdown IE, 5 males, 5 females). B) Active and Inactive lever presses for mice prior to injection of the control LacZ gRNA or knockdown IE gRNA AAV. Both groups demonstrated preference for the active lever, and there were no differences in the active or inactive lever responding between groups. C) Mice injected into the VTA with the control LacZ gRNA exhibited a robust incubation of craving response, with greater active lever presses on day 21 (AD21) compared to day 1 (D1) of forced abstinence (n=12; 7 males, 5 females). D) Mice injected into the VTA with the knockdown IE gRNA did not exhibit incubation of cocaine craving (n=10; 5 males, 5 females). E) In comparing the percent change in active lever presses from day 21 to day 1, the control LacZ and IE gRNA groups exhibited opposing responses (n=5-7/group; control, 7 males, 5 females; knockdown IE, 5 males, 5 females). F) No differences were found in the number of food pellets earned by control LacZ gRNA and knockdown IE gRNA mice during baseline (B1-3) and reversal sessions (R1-4) of the food lever reversal task (n=9-11/group; control, 6 males, 5 females; knockdown IE, 5 males, 4 females). G) Number of active and inactive lever presses in control and knockdown mice during the food lever reversal task. Statistics: ns = not significant, **p<0.01, ***p<0.001, ****p<0.0001. Error bars indicate standard error mean (SEM).

For the *lacZ* control versus *Bdnf* P1 cohort, prior to AAV injection on AD1, mice that were pre-assigned to different AAV groups had indistinguishable levels of responding for the total number of infusions (**Fig. 4A**; Unpaired t-test, t_(30)_=0.8764, p= 0.3878). Both groups distinguished between the active and inactive levers, and they did not differ in the number of active or inactive lever presses (**Fig. 4B**; Two-way Repeated Measures (RM) ANOVA, Group F_(1, 30)_= 0.3550, p=0.5558; Lever F_(1, 30)_=129.0, p<0.0001; Interaction F_(1, 30)_=0.1425, p=0.7085; Tukey’s post-hoc, active vs inactive: control p<0.0001, P1 knockdown p<0.0001). When cocaine seeking behaviors at AD1 and AD21 were compared, an increase in active lever presses was observed in the control *lacZ* gRNA group, indicating an incubation of cocaine seeking (**Fig 4C**; Paired t-test, t_(15)_=4.727, p= 0.0003). By contrast, incubation was not seen in the *Bdnf* P1 gRNA group, and in fact, there was a significant decrease in active lever presses between AD1 and AD21 (**Fig 4D**; Paired t-test, t_(15)_=3.320, p= 0.0047). These data are also represented as a percent change in active lever presses between AD1 and AD21, where the *Bdnf* P1 gRNA group is significantly lower than the *lacZ* control (**Fig 4E**; t-test, t_(30)_=4.792, p<0.0001).

To verify that the reduction in lever pressing was due to cocaine-related responding and not secondary to the animal’s ability to press the lever or reward learning in general, subjects were then trained in the food lever reversal task. Subjects acquired stable responding for food pellet rewards during three baseline sessions (B1-B3). The lever assignment was then switched on reversal day 1 (R1), and mice were maintained with the new lever assignment across an additional 3 days (R2-R4). When examining the number of food pellets earned, there were no significant differences between groups across sessions, either during baseline responding or during the reversal sessions (**Fig 4F**; Two-way RM ANOVA, Group F_(1, 11)_=3.503, p=0.0881; Session F_(6, 66)_=8.564, p<0.0001; Interaction F_(6, 66)_=1.693, p=0.1365). In examining the active and inactive lever responding, the knockdown *Bdnf* P1 gRNA group exhibited a lower perseverance of responding for the inactive lever on the first reversal day compared to the control *lacZ* gRNA group, but both groups did not differ with responding on the active lever across all sessions (**Fig 4G**; Two-way RM ANOVA, Group F_(3, 22)_=31.85, p<0.0001; Session F_(6, 132)_=6.976, p<0.0001; Interaction F_(18, 132)_= 12.43, p<0.0001; Tukey’s post-hoc, R1 session, control inactive vs P1 knockdown inactive p=0.0009). These data verify that *Bdnf* knockdown in the VTA specifically affects incubation of cocaine seeking during abstinence and does not cause a general deficit in the ability of the mice to press a lever with a high level of effort to earn a non-drug reward.

We saw similar results in an independent conditional dCas9-KRAB cohort when we compared the *lacZ* control gRNA to the *Bdnf* IE gRNA. Again, prior to viral injection both assigned groups showed no difference in responding for the total number of cocaine infusions (**Fig. 5A**; Unpaired t-test, t_(20)_=0.3452, p=0.7336) and both distinguished between the active and inactive levers (**Fig. 5B**; Two-Way RM ANOVA, Group F_(1, 20)_=0.0436, p=0.8367; Session F_(1, 20)_=40.77, p<0.0001; Interaction F_(1, 20)_=0.2146, p=0.6482; Tukey’s post-hoc, active vs inactive: control p=0.0003, IE knockdown p=0.0002). The *lacZ* control gRNA group showed significantly increased active lever pressing from AD1 to AD21 (**Fig. 5C**; Paired t-test, t_(11)_=3.619, p=0.0040), but the *Bdnf* IE gRNA group did not (**Fig. 5D**; Paired t-test, t_(9)_=1.649, p=0.1336). As a result the percent change in active lever presses between AD1 and AD21 was significantly lower in the *Bdnf* IE gRNA group compared with the *lacZ* control (**Fig. 5E**; Unpaired t-test, t_(20)_=3.031, p=0.0066). When the mice were trained in the food lever reversal task, a significant decrease in the number of food pellets earned was found on the first reversal session, with the IE knockdown group showing a deficit compared to the control group (**Fig. 5F**; Two-way RM ANOVA, Group F_(1, 20)_=3.655, p=0.0703; Session F_(6, 120)_=11.07, p<0.0001; Interaction F_(6, 120)_=3.927, p=0.0013; Tukey’s post-hoc, R1 session, control vs IE knockdown p<0.0001). This difference was also observed in the number of active lever presses on the first reversal session (**Fig. 5G**; Two-way RM ANOVA, Group F_(3, 40)_=60.36, p<0.0001; Session F_(6, 240)_=28.75, p<0.0001; Interaction F_(18, 240)_=16.83, p<0.0001; Tukey’s post-hoc, R1 session, control active vs IE knockdown active p<0.0001). However, no differences were found in subsequent reversal sessions, indicating a transitory effect on cognitive flexibility with knockdown of the IE *Bdnf* region; this is in contrast to knockdown at promotor I in which enhanced cognitive flexibility was observed with a decrease in the perseverance of responding on the inactive lever at this time point. These findings indicate minor differences may be present with knockdown strategies that include *BdnfI* compared to *BdnfI* and *BdnfIV* transcripts. Overall, these data demonstrate that local inhibition of *Bdnf* induction in the VTA during abstinence, either by repressing promoter I or the intragenic *Bdnf* enhancer, is sufficient to prevent incubation of cocaine seeking.

### Identification of a program of gene transcription induced in VTA neurons by cocaine abstinence

To identify genes in addition to *Bdnf* that undergo transcriptional regulation during abstinence after chronic cocaine, and that might also contribute to drug seeking during abstinence, we isolated the nuclei of VTA neurons for sequencing. We genetically tagged the nuclei of DAT-Cre expressing neurons by crossing these mice to a strain expressing a Cre-inducible transgene of the nuclear envelope protein Sun1 fused on the cytoplasmic side with GFP (Gallegos et al., 2023; Mo et al., 2015) (**Fig. S5A)**. We confirmed that nuclei labeled by the Sun1-GFP transgene colocalized with tyrosine hydroxylase (TH) immunostaining in the VTA (**Fig S5B**). Following 7d of repeated saline or cocaine (20mg/kg, i.p.) injections, brains were harvested on AD1 or AD14 for VTA microdissection. GFP tagged nuclei were enriched by Miltenyi MACS anti-GFP Microbeads for single nucleus RNA-sequencing (snRNAseq) or by anti-GFP coated Dynabeads for bulk RNA-seq and chromatin sequencing. DAT neuron enrichment in the immunoprecipitated fraction was confirmed by visualization of GFP+ nuclei in the isolated fractions and by enrichment of *Th* and *Slc6a3* (**Fig. S5C-E**). Notably, although both protocols enriched for DA neurons relative to the input sample, the Miltenyi MACS purification still included many GFP-nuclei, which we used to our advantage in our single nucleus sequencing study.

We conducted snRNAseq with the Scale Bioscience ScalePlex kit, which allows us to track and combine isolated neurons from each single mouse in one sequencing experiment. We used 4-6 adult, male and female mice from each of the following four conditions: saline AD1, saline AD14, cocaine AD1, cocaine AD14. After QC we retained 46,612 high quality nuclei spread equally across the four treatment groups (**Fig S6A-B)**. Following data integration, unsupervised dimensionality reduction revealed 19 transcriptionally distinct clusters **(Fig 6A**). Clusters 3 and 7 contained 5891 nuclei that showed enriched expression of *Th* (encoding for tyrosine hydroxylase), *Slc6a3* (encoding for the dopamine transporter), *Slc18a2* (encoding for vesicular monoamine transporter), and *Ddc* (encoding for dopamine decarboxylase), identifying them as the dopaminergic neuron clusters (**Fig 6B**). To verify the identity of these cells, we utilized a published VTA snRNAseq dataset (Phillips et al., 2022) and transferred the anchors onto our snRNAseq dataset (**Fig S6C**). This comparison reconfirmed our designation of clusters 3 and 7 as VTA DA neurons. Prior studies have also identified transcriptional subtypes of DA neurons that differ in their anatomical localization, function, and circuit connectivity (Poulin et al., 2018; Poulin et al., 2020; Poulin et al., 2014). Subsetting and dimensionality reduction of our DA neurons revealed 7 transcriptionally distinct clusters (**Fig S6D,E**), with nuclei from all four treatment groups spread equally across all clusters (**Fig S6F**). We categorize our subcluster 0 as DA^1B^ (*Aldh1a1^-^, Sox6^+^, Ndnf ^+^, Tacr3^+^*), subcluster 1 as DA^2D^ (*Aldh1a1^-^, Sox6^-^, Calb1^+^, Vip^+^, Cck^+^*), subcluster 2 as DA^2A^ (*Calb^+^, Slc17a6^+^, Adcyap1^+^*), subcluster 3 as DA^1A^ (*Aldh1a1^+^, Sox6^+^, Ndnf^+^*), and subcluster 6 as DA^2B^ (*Lpl^+^, Grp^+^, Aldh1a1^+^, Calb1^+^, Tacr3^+^*) (**Fig S6E**).

**Figure 6:**
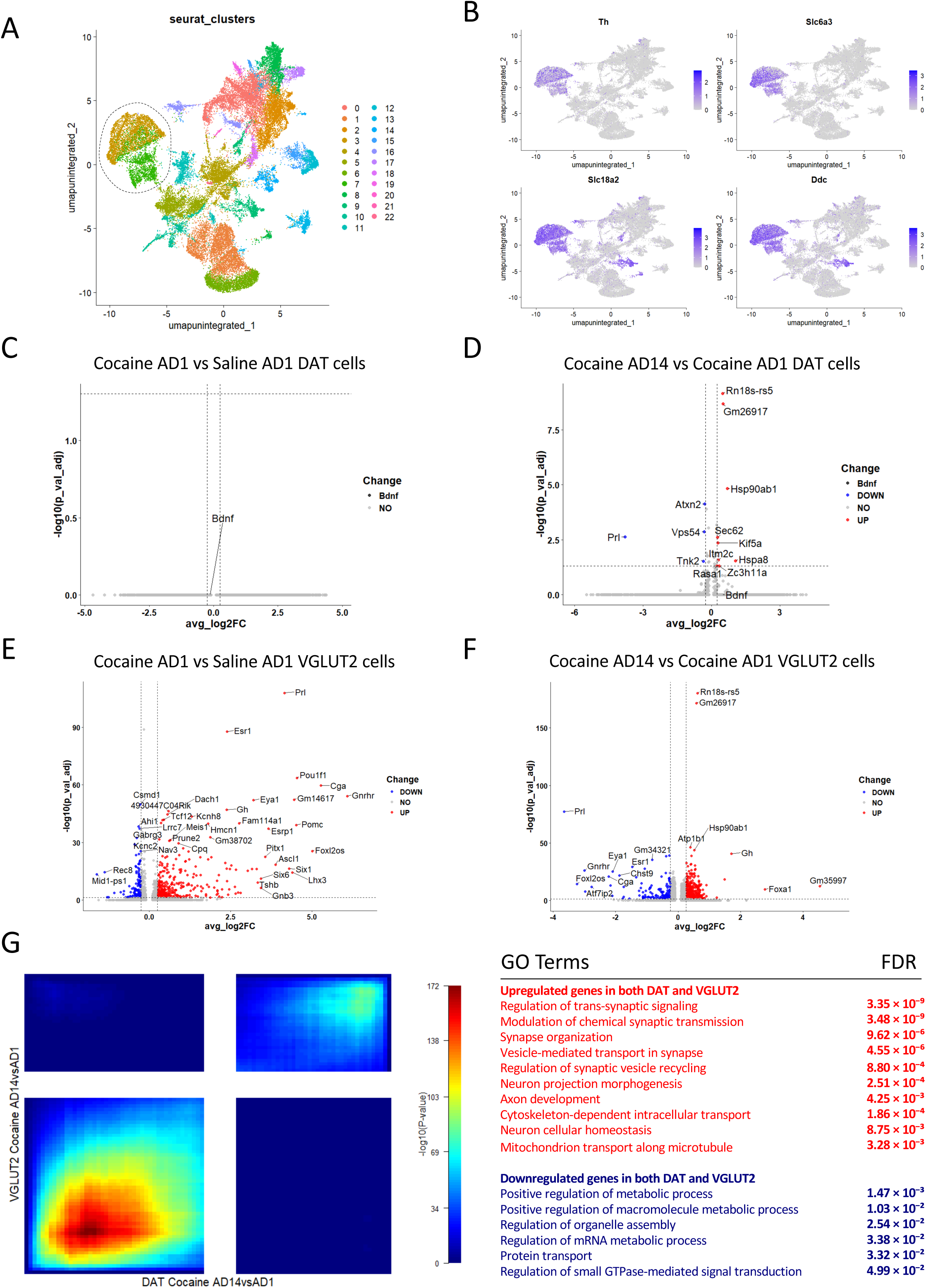
Identification of a gene expression program induced in VTA DAT and VGLUT2 neurons by abstinence after cocaine. **A)** UMAP of cell types from the GFP enriched fraction of VTAs from saline/cocaine treated mice. **B)** Feature plots for classic dopaminergic genes Th, Slc6a3, Slc18a2, and Ddc, identifying the dopaminergic cluster. **C)** MAST analysis determining differentially expressed genes in all dopaminergic cells between cocaine and saline at AD1 conditions. Average log2FC cut off set to ±0.25 (20% difference). **D)** MAST analysis determining differentially expressed genes in all dopaminergic cells between cocaine AD14 and AD1 conditions. Average log2FC cut off set to ±0.25 (20% difference). **E)** MAST analysis (above) determining differentially expressed genes in all glutamatergic cells between cocaine and saline at AD1 conditions. FDR <0.05 and |L2FC|>0.25. **F)** MAST analysis (above) determining differentially expressed genes in all glutamatergic cells between cocaine AD14 and AD1 conditions. FDR <0.05 and |L2FC|>0.25. **G)** Rank-rank hypergeometric overlap (RRHO) plot displaying concordance of differential gene expression measured by RNA-seq in Glutamatergic cells between cocaine AD14 and AD1 (y-axis), and in Dopaminergic cells between cocaine AD14 and AD1 (x-axis). GO terms for the common differential genes (right).

To identify VTA DAT neuron genes that show differential regulation during cocaine abstinence, we used MAST to perform zero-inflated regression analysis by fitting a linear mixed model (LMM) (**Fig 6C**) (Finak et al., 2015). When we compared DAT neuron gene expression on AD1 between mice that received cocaine and those that received saline we saw no differentially regulated genes, suggesting that repeated cocaine exposure alone does not cause transcriptional adaptations in VTA DA neurons. By contrast, when we compared AD1 and AD14 nuclei from mice that received cocaine, we identified 9 upregulated genes and 4 downregulated genes at an FDR corrected p value<0.05 and a |log_2_fold change| >0.25 (**Table S3**). Of note, although *Bdnf* passed the fold change threshold, it did not pass the significance threshold, likely because of its very low expression in DAT neuron nuclei relative to the genes called significant. Together these data show that DAT neurons experience dynamic transcriptional adaptations selectively during the period of abstinence following repeated cocaine exposure.

In addition to the VTA DAT cells, our snRNAseq dataset contained a large number of glutamatergic nuclei that again had all four treatment groups spread equally across all clusters (**Fig S6G,H**). To identify differentially expressed genes in glutamatergic neurons in response to either cocaine or abstinence, we performed MAST on these clusters (**Fig 6E,F; Table S4**). Unlike our the VTA DAT cells, the glutamatergic cells showed dynamic gene expression changes at AD1 when we compared nuclei from mice exposed to saline with those exposed to cocaine (**Fig 6E**). Glutamatergic cells also showed differential gene expression during abstinence, when we compared samples at AD14 to AD1 in mice exposure to cocaine (**Fig 6F**). A Rank-Rank Hypergeometric Overlap (RRHO) analysis comparing the abstinence-dependent gene programs induced in the DAT and VGLUT2 cells showed a strong overlap in the upregulated genes between both cell types, and a much lower overlap in the downregulated genes (**Fig 6G**). Gene Ontology analyses on these genes showed an enrichment of signaling and synaptic terms, such as regulation of trans-synaptic signaling (*Slc17a6, Slc6a3, Gabbr1, Cplx1*, and *Syn1*), modulation of chemical synaptic transmission (*Prkaca, Plcg1, Pde2a, Camk2n1*, and *Grm3*), and synapse organization (*Lrrtm2, Pcdh10, Cntnap1, Celsr2, Flrt3,* and *Sez6l2*), while the terms for the downregulated genes showed an enrichment for broader regulatory processes, such as metabolic process (*Rptor, Adk, Pank2, Ip6k2,* and *Pex14*) and organelle assembly (*Stx18, Vps54, Exoc5, Nup98*, and *Rab11fip3*) (**Table S5**). The strong overlap in differentially upregulated genes between VGLUT2 cells and DAT cells suggests that there may exist a common cocaine abstinence program induced in both cell types that modulates synaptic signaling in the VTA.

### Egr1 as an upstream regulator of abstinence-induced gene expression in VTA neurons

Our snRNAseq data identified a set of genes beyond *Bdnf* that are differentially regulated in VTA neurons during a weeks-long period of abstinence after cocaine. These data raise the question of the transcriptional regulatory mechanisms that coordinate this abstinence program of gene expression. The cell-type specific actions of transcription factors are determined by chromatin features that control the accessibility and 3D contacts promoters and enhancers in the genome (Griffith et al., 2024). Thus to discover abstinence-relevant transcriptional mechanisms, we characterized the chromatin architecture of DA neuronal nuclei isolated from the VTA of mice using the highly efficient Dynabeads-INTACT protocol shown in **Fig. S5**.

We used a novel low-input chromatin conformation capture method called Hi-C at accessible DNA regulatory elements (HiCAR) (Wei et al., 2022). This method relies on the transposase Tn5 to capture 3D genome interactions with regions of accessible chromatin and therefore is optimal for determining the promoter-enhancer regulatory architecture of cells (Park et al., 2025; Wei et al., 2026). To validate the sensitivity of HiCAR to identify cell-type specific chromatin features in neuronal nuclei purified from the adult mouse brain, we tested the ability of HiCAR to identify transcriptionally relevant differences in chromatin structure between DA neurons isolated from the VTA with cerebellar granule neurons (CGNs) (**Fig. S7**). Quantitative comparison of these HiCAR datasets with RNA-seq data identified thousands of cell-type specific accessible regions and chromatin loops that correlate with differential gene expression **(Fig. S7A-C).** Specificity of the accessibility and looping data from the HiCAR pipeline can be seen at genes like *Gabra6*, which encodes a GABA_A_ receptor subunit that is selectively expressed in CGNs, and *Slc6a3*, encoding the dopamine transporter, which is a canonical marker of DA neurons **(Fig S7D-F)**.

To determine if there were changes in chromatin accessibility or promoter-enhancer looping induced in VTA DA neurons during cocaine abstinence, we made HiCAR libraries from VTA DA neurons isolated following 14d of abstinence after cocaine exposure and compared them to libraries we made from saline exposed mice as control. However, we found no significant differences genome-wide in the pattern of accessible chromatin between treatment conditions in DA neurons **(Fig S8A)**, and no significant differences in chromatin looping in DA neurons isolated from the saline or cocaine treated mice **(Fig S8B)**. These data suggest that, rather than changing chromatin architecture, prior exposure to cocaine may cause DA neurons to reconfigure the transcriptional control of their existing chromatin regulatory elements to adapt gene expression programs over the course of abstinence.

To discover possible transcription factor (TF) regulators of the cocaine abstinence program in VTA DA neurons, we used our DA neuron-specific HiCAR data to define the experimentally validated topologically associating domains (TADs) containing the abstinence-regulated genes from our snRNA-seq study. Enhancers most commonly regulate genes within the same TAD (Pachano et al., 2022), thus these data allowed us to narrow our search for TF binding sites to the most functionally-relevant enhancers. Using the sequence of all accessible chromatin in DA neurons for comparison, we applied the motif discovery program HOMER to identify transcription factor (TF) binding motifs enriched within accessible chromatin of the TADs containing our abstinence-regulated genes from DA neurons **(Fig 7A)**. This analysis identified the Egr1 TF binding motif to be enriched in the open chromatin sequences in the TADs of our DEGs. Prior studies in mice have reported significantly increased *Egr1* mRNA expression in the VTA 1hr after cocaine IVSA (Campbell et al., 2021) or weakly increased mRNA levels 24hr after IVSA (Carpenter et al., 2020). *EGR1* mRNA was also found to be elevated in the ventral midbrain in a postmortem study of human cocaine users relative to age-matched controls, and the same study showed that EGR1 protein prominently localized to DA neurons in human VTA (Bannon et al., 2014). To determine if Egr1 protein was regulated by cocaine exposure in the VTA of our mice, we used mice from our four treatments groups (7d cocaine or saline followed by 1 or 14 days of abstinence) for Egr1immunofluorescence. We observed significantly increased Egr1 expression in the VTA of mice 1 day after repeated cocaine exposure relative to saline exposed controls (**Fig 7B-D)**. These Egr1 positive cells were found to colocalize with TH expression (**Fig S9**), suggesting they are DAT neurons. The elevated expression of Egr1 did not persist, however, as it was not different in cocaine exposed mice after 14d of abstinence compared with their saline controls. These data suggest that transient Egr1 induction may prime genes including *Bdnf* for later regulation during abstinence, an idea we discuss further below.

**Figure 7:**
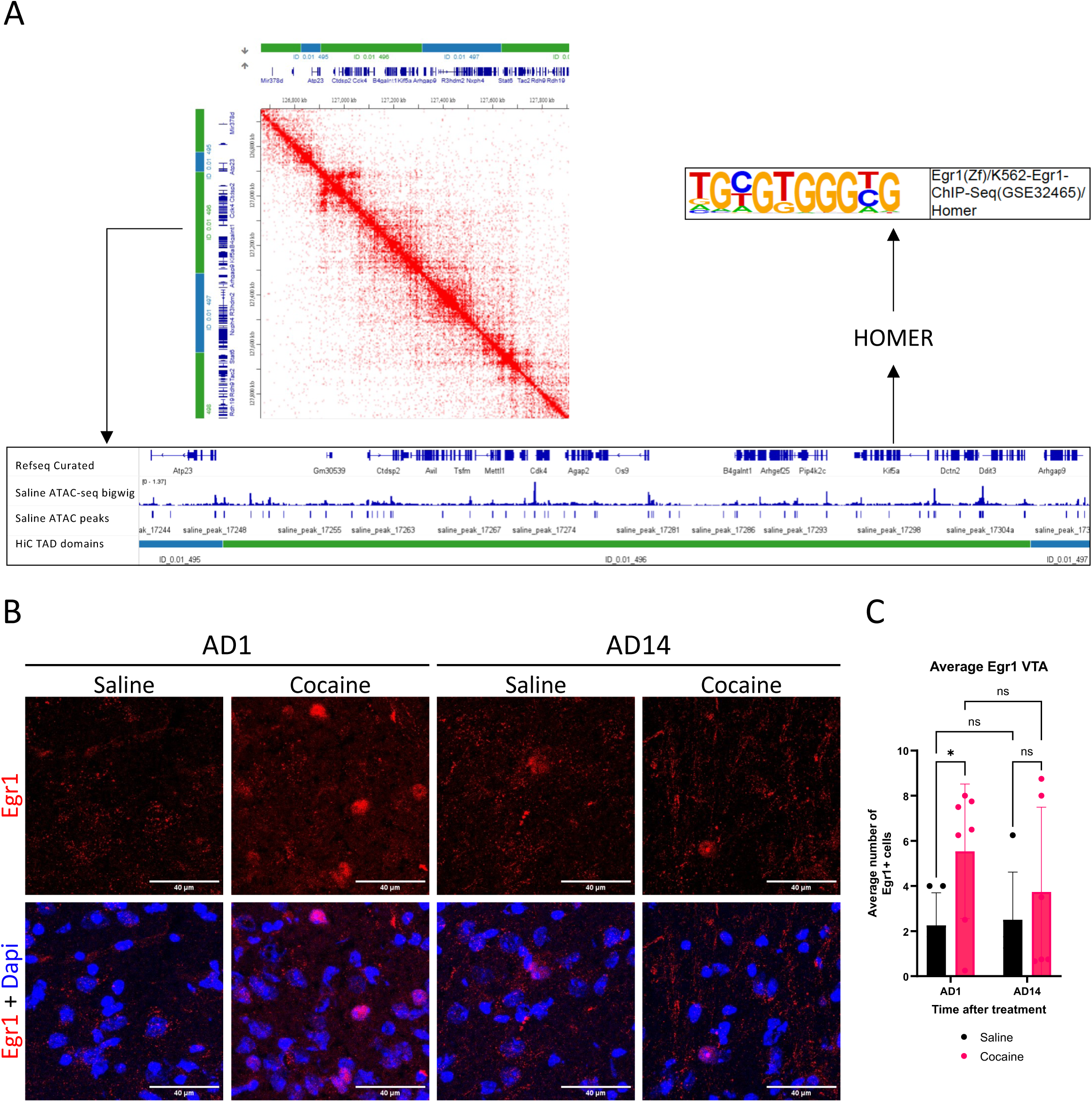
Identification of Egr1 as a candidate regulator of DAT abstinence-regulated genes. **A)** Overview of the workflow for mapping ATAC-seq peaks to TADs of differentially regulated genes and identifying enriched Egr1 motifs using HOMER. **B)** Representative immunohistochemistry images of Egr1 signal (red) in the medial VTA of mice treated with saline or cocaine at AD1 (left panels) and at AD14 (right panels). **C)** Average Egr1 signal in the VTA in each animal upon cocaine abstinence. Statistics: n=5-7mice, 50-70 cells sampled per animal, 8C: Two-way ANOVA with post-hoc Uncorrected Fisher’s LSD; ns = not significant, *p<0.05. Error bars indicate standard deviation (SD).

## Discussion

Here we show that abstinence-induced expression of *Bdnf* in DAT neurons is mediated by *Bdnf* promoter I as well as an intragenic enhancer, and we find that CRISPR inhibition of *Bdnf* expression via either regulatory element is sufficient to block incubation of cocaine seeking in mice. These data indicate that the induction of *Bdnf* in the VTA during abstinence plays a causal role in cocaine seeking behaviors, demonstrating the functional role of VTA transcriptional plasticity in cocaine seeking behavior. In snRNA-seq data, we identify a convergent program of gene expression induced in DAT and VGLUT2 neurons of the VTA selectively by cocaine abstinence, broadening understanding of this plasticity process. We propose that changes in VTA neuron function during abstinence that result from these transcriptional changes may contribute to the risk of relapse in cocaine use disorder.

### Distinct regulatory elements govern cell-type specific Bdnf transcription in the VTA

BDNF is a well-established plasticity effector that plays important roles in coupling experience-dependent synaptic signaling with changes in synapse strength and circuit connectivity in many different cell types and contexts (Park & Poo, 2013). Transcriptional control is the major mechanism used by neurons to regulate BDNF availability, thus study of this gene offers a functionally important window into the stimulus-regulated cellular transcriptional mechanisms that coordinate circuit plasticity (West et al., 2014). An important conceptual takeaway from our study is that the complex regulation of *Bdnf* transcription offers a means to control the cell-type specific actions of this factor. *Bdnf* is regulated at the transcriptional level by at least eight different promoters as well as both intragenic and distal enhancer elements (Brookes et al., 2023; Timmusk et al., 1994; Tuvikene et al., 2021). Although all of the transcripts produced by the *Bdnf* locus encode the same mature BDNF protein, there has long been evidence that distinct *Bdnf* transcripts are expressed under different conditions (West et al., 2014). Each of the regulatory elements of the *Bdnf* gene is controlled by different complements of transcription factors, providing a mechanism to selectively couple *Bdnf* induction in a given cell type or to different upstream stimuli.

Prior studies found a selective increase in *Bdnf* mRNA made from promoter I in the VTA during cocaine abstinence (Schmidt et al., 2012), suggesting that the accumulation of BDNF protein observed in this brain region during cocaine abstinence might be due to local activation of this promoter (Grimm et al., 2003). Consistent with this hypothesis, we show that CRISPR inhibition of *Bdnf* promoter I blocks the abstinence-induced elevation of *Bdnf* mRNA in VTA. Furthermore, we show that although *Bdnf* is expressed in both DAT and VGLUT2 expressing neurons in the VTA, the induction of *Bdnf* during abstinence is selective for DAT neurons. Inhibiting promoter I only leads to a significant reduction of *Bdnf* in the DAT neurons but not the VGLUT2 neurons, supporting that different cell types in the VTA control *Bdnf* expression through distinct transcriptional mechanisms. Interestingly, dCas9-KRAB recruitment to the intragenic enhancer represses *Bdnf* expression in both DAT and VGLUT2 cells, suggesting that this enhancer can cooperate with either the upstream (promoter I-III) or downstream (promoter IV-VII) promoter clusters in the *Bdnf* gene through 3D chromatin folding (Beagan et al., 2020). These detailed data on the cell-type specificity of *Bdnf* regulation in the VTA complement studies showing that different cell-types respond to common stimuli with distinct patterns of gene expression changes (Gallegos et al., 2023; Whitney et al., 2014). This non-uniform plasticity allows different cell types to adapt in distinct ways, supporting circuit-level adaptations that contribute to behavioral changes (Griffith et al., 2024).

### Consequences of VTA BDNF synthesis for circuit plasticity underlying cocaine seeking

Our data on the reduced incubation of cocaine seeking following local CRISPR/dCas9-KRAB inhibition of *Bdnf* in the VTA provide direct evidence of a requirement for abstinence-induced VTA *Bdnf* synthesis in the incubation of cocaine seeking during forced abstinence. These data both complement and extend prior studies showing that infusion of BDNF into the VTA is sufficient to promote cocaine seeking behaviors (Grimm et al., 2003; Lu et al., 2004). Notably, the VTA is not the only brain region where functionally important transcriptional adaptations occur during cocaine abstinence that contribute to the incubation of seeking behaviors. Significant changes in gene expression have been profiled in D1 DA receptor expressing medium spiny neurons in the NAc (Kawa et al., 2025) as well as in prelimbic cortex (Barry et al., 2026) of rats after various times of abstinence following cocaine self-administration. Furthermore, one study showed that in the ventral hippocampus, induction of the calcium buffering protein calretinin by FosB/ΔFosB in NAc-projecting neurons causes changes in excitability and is necessary for cocaine reward (Eagle et al., 2026).

These data raise the importance of thinking about how the regulation of genes in one part of the circuitry that underlies cocaine seeking can impact the functional output of the network. BDNF is a secreted protein, thus it provides a useful example for understanding how intrinsic changes in gene expression alter circuit connectivity. BDNF protein can be transported in both an anterograde (Altar et al., 1997) and a retrograde manner (Watson et al., 1999) to impact neurons presynaptic or postsynaptic to its source of synthesis. In the context of its induction during cocaine abstinence, BDNF protein synthesized in the VTA could be trafficked anterogradely for release from DAT neuron terminals in the NAc. This would trigger aspects of cocaine seeking that require activation of the BDNF receptor TrkB in the NAc (Li et al., 2013). In parallel, there is also evidence for actions of BDNF in the VTA through paracrine signaling to the inputs on DA neurons. A prior study has observed that upon cocaine abstinence, excitatory synapses onto the VTA DA neurons were susceptible to potentiation in a BDNF-TrkB manner (Pu et al., 2006). BDNF-dependent strengthening of excitatory synaptic connections onto dopaminergic neurons has the potential to augment transfer of salient cocaine-related information.

### Chromatin-based regulation may underlie persistent transcriptional plasticity during cocaine abstinence

Our snRNAseq data reveal that the drug-free period after chronic cocaine exposure is associated with significant transcriptional plasticity in VTA neurons that extends well beyond *Bdnf*. Prior bulk RNA-seq studies have reported transcriptional changes in VTA after home cage or IVSA cocaine, however they could not deconvolve which cells showed these changes (Campbell et al., 2021; Carpenter et al., 2020). Our data reveal that some of the transcriptional responses to cocaine are cell-type specific. For example, one day after repeated cocaine we observed a significant transcriptional response in VGLUT2+ neurons compared with mice that received saline, whereas we detected no significant transcriptional changes in the DAT+ neurons of the same mice. This may be a consequence of the small population of DAT neurons recovered even though we enriched these cells about 5X above their detection in an unbiased VTA snRNAseq study (Phillips et al., 2022). Furthermore the sparsity of snRNA-seq data limits our ability to detect lowly expressed genes; for example, we were unable to detect induction of *Bdnf* in the snRNAseq data even though we quantitatively validated induction of this gene by RNAscope FISH. Despite these limitations, we identified significant gene expression changes in both DAT-and VGLUT2-expressing neurons after 14 days of cocaine abstinence, and when we performed a rank-rank hypergeometric overlap comparison we found that there were common biological processes induced during abstinence in both cell populations. Upregulated genes were significantly associated with synapse terms, whereas downregulated genes belonged to broader metabolic categories. These data are consistent with a plasticity of connectivity in the VTA during abstinence, which may underlie future cocaine seeking behaviors when the mouse is returned to the IVSA chamber.

It is striking that the gene expression changes arising during abstinence are both slow to begin after cessation of cocaine exposure and persistent (days-weeks) once induced. This time course suggests a very different mechanism of transcriptional regulation compared with the rapid and transient phosphorylation of stimulus inducible transcription factors such as CREB that underlie the rapid (minutes-hours) responses to cocaine (Carlezon et al., 1998). Epigenetic modifications of chromatin can underlie very long-lasting differences in gene expression and have been proposed to be targets of regulation by drugs of abuse including cocaine. Consistent with this possibility, others have reported an increase of acetylation of histone H3 at *Bdnf* promoter I (Schmidt et al., 2012) as well as an accumulation of dopaminylated histone H3 in the VTA during cocaine abstinence (Lepack et al., 2020). We considered the possibility that abstinence-induced changes in chromatin architecture might reconfigure promoter-enhancer interactions to alter gene expression levels. We used a chromatin conformation method called HiCAR that leverages the transpose Tn5 to detect both open chromatin and chromatin architecture in order to enrich for regulatory interactions, and we confirmed that this method can reveal cell-type specific differences in chromatin architecture in neuronal nuclei isolated from the adult mouse brain (Wei et al., 2026). However, we detected no significant abstinence-induced changes in a purified population of VTA DA neurons. A limitation is that we performed this assay in bulk, so if only a small subset of the neurons were responding we might have obscured the change.

However, we were able to leverage this information to discover transcription factor binding sites that are enriched in the chromatin regions surrounding our abstinence regulated genes. These data suggested a role for the immediate-early gene (IEG) transcription factor Egr1, which is known to be induced in the VTA 24 hr after cessation of IVSA cocaine (Campbell et al., 2021) and is an important upstream regulator of *Bdnf* transcription (Esvald et al., 2025; Tuvikene et al., 2021). We confirmed that Egr1 expression is found in DAT neurons of the VTA 24 hr after cessation of chronic cocaine; however we saw no expression at 14 days of abstinence when our transcriptional program is active. Interestingly, prior studies have suggested that transient induction of IEG transcription factors can prime genes for later regulation in the context of learning and memory (Marco et al., 2020) as well as across stages of the estrus cycle in female mice (Rocks et al., 2025). Future studies that target Egr1 directly will help to resolve the functional importance of this transcription factor in the context of abstinence-induced transcriptional memory in VTA neurons.

## Acknowledgements

This work was supported by NIH grants R21DA061547 (A.E.W and C.D.F) and F31NS127573 (A.N.). We thank Tiffany Ko, Aryana Yousefzadeh, and Xiaolu Yang for technical assistance, and the Duke University Mouse Behavioral and Endocrine Analysis Core facility for the use of equipment and experimental support.

