## Supplementary material for "Transcriptional plasticity of ventral tegmental area neurons induced after cessation of chronic cocaine exposure is required for incubation of cocaine seeking": Figures S1-S9

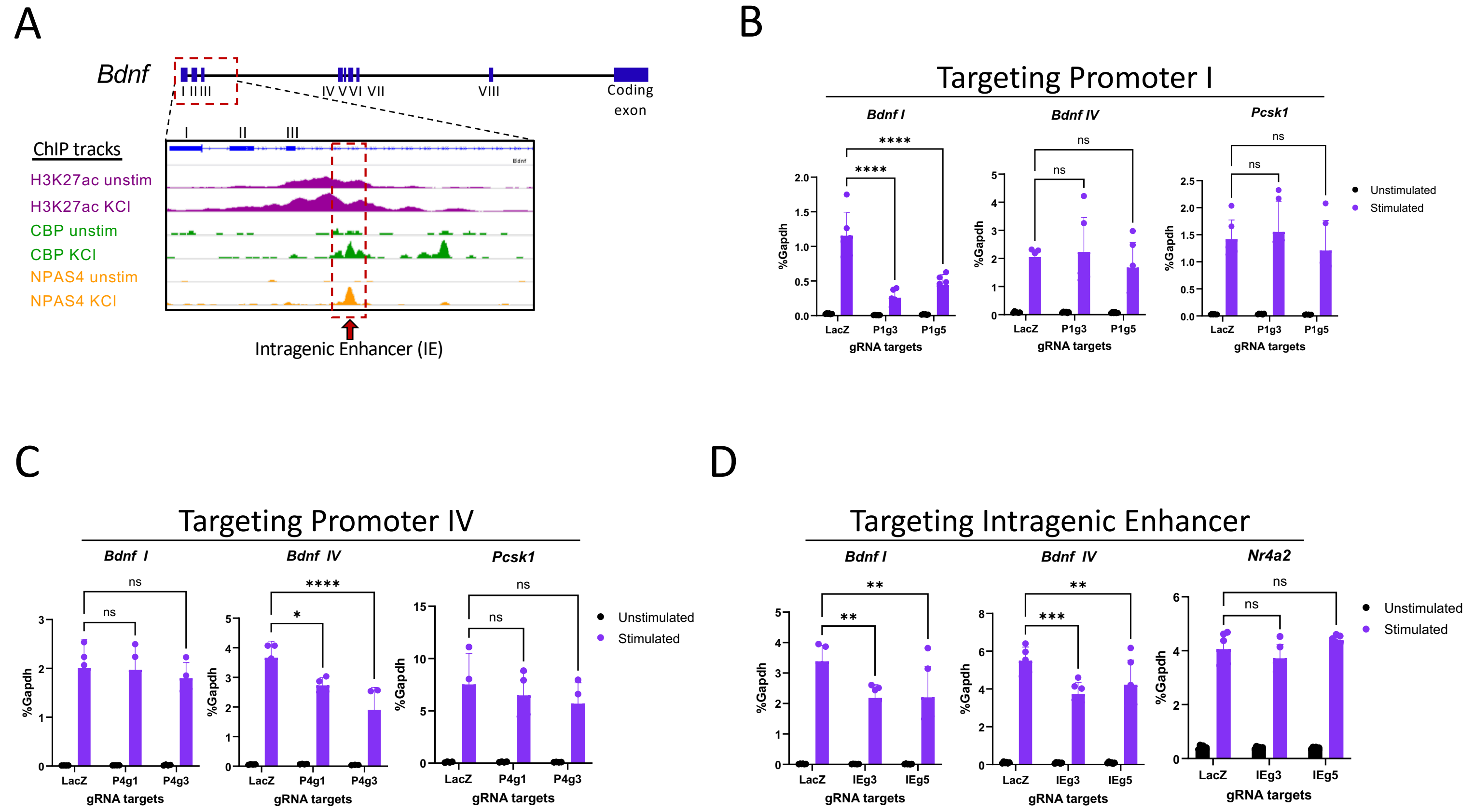


**Supplemental Figure S1: dCas9-KRAB targeting different regulatory elements can repress *Bdnf* in culture**

A) Representative image of *Bdnf* genomic locus with 8 non-coding exons and one non-coding exon. Additionally, there is an activity-dependent intragenic enhancer located downstream of exon III, demarcated by the red box showcasing the increase in chromatin features such as H3K27ac, CBP, and NPAS4 binding upon KCl stimulation (Kim et al., 2010).

B) RT-qPCR data from unstimulated and KCl stimulated cortical neuron cultures infected with dCas9-KRAB and gRNAs targeting promoter I.

C) RT-qPCR data from unstimulated and KCl stimulated cortical neuron cultures infected with dCas9-KRAB and gRNAs targeting promoter IV.

D) RT-qPCR data from unstimulated and KCl stimulated cortical neuron cultures infected with dCas9-KRAB and gRNAs targeting the intragenic enhancer.

Statistics: n=4-6 (2-3 independent experiments with 2 biological replicates per experiment), Two-way ANOVA with post-hoc Tukey's multiple comparison; ns = not significant, *$p<0.05$, **$p<0.01$, ***$p<0.001$, ****$p<0.0001$

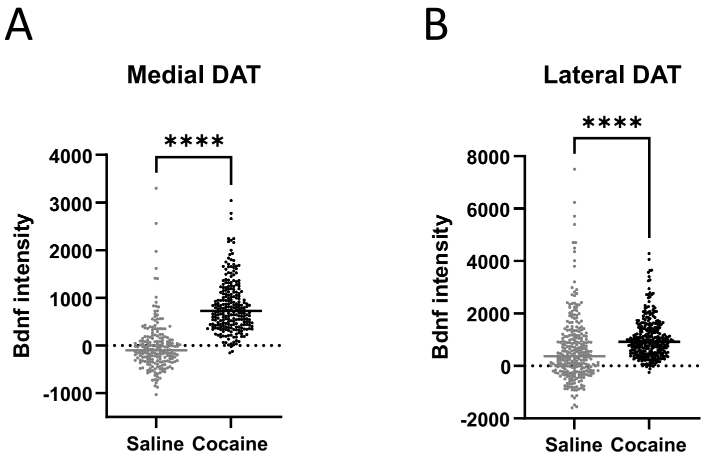


**Supplemental Figure S2: *Bdnf* upregulation upon cocaine abstinence is seen in VTA DAT cells in mice injected with AAV.**

A) Quantification of *Bdnf* signal in medial DAT cells from mice injected with LacZ gRNA AAV.

B) Quantification of *Bdnf* signal in lateral DAT cells from mice injected with LacZ gRNA AAV.

Statistics: n=4 mice, 50-70 cells sampled per animal, Unpaired t-test, ****p<0.0001

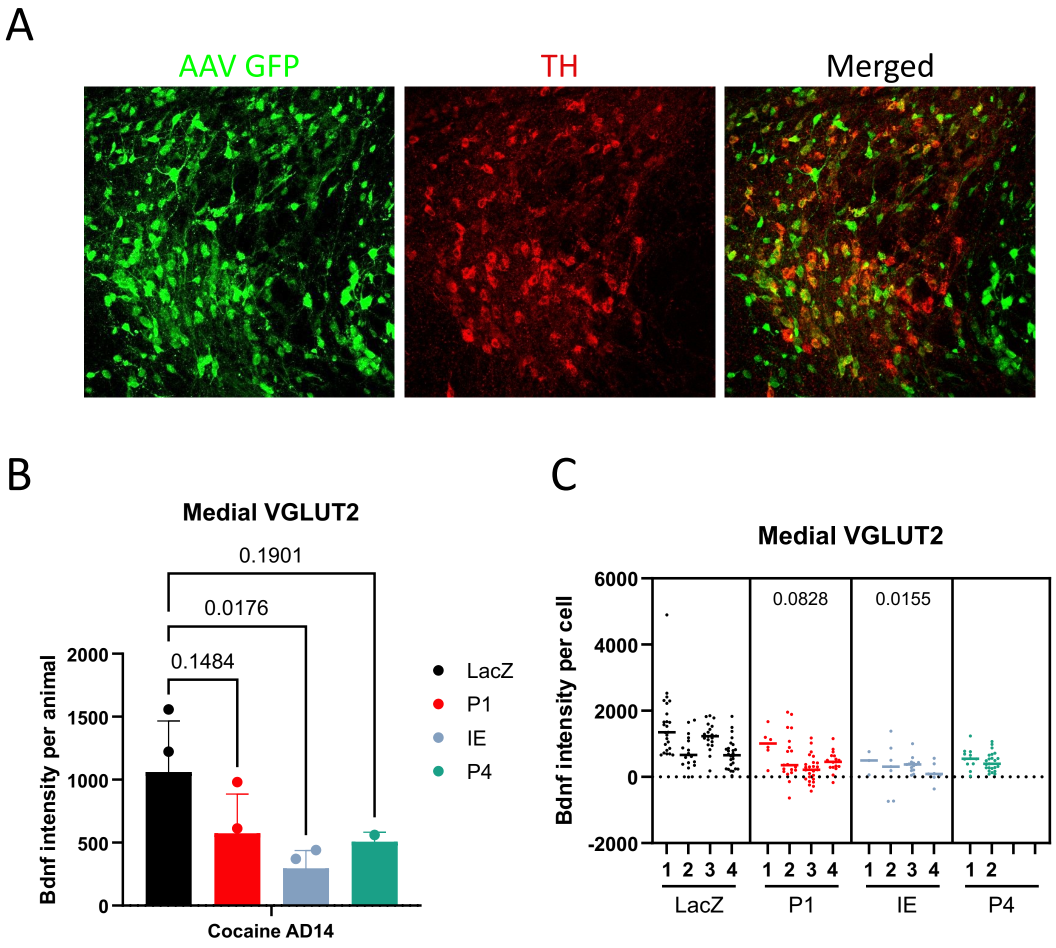


**Supplemental Figure S3: *Bdnf* upregulation upon cocaine in VTA VGLUT2 cells**

A) Representative immunohistochemistry images of GFP from AAV (medial) in TH+ (red) and TH- cells in the VTA of mice injected with LacZ gRNA AAV.

B) Quantification of *Bdnf* signal in medial VGLUT2 cells as average *Bdnf* signal per mouse injected with LacZ, P1, IE, or P4 gRNA AAV, treated with cocaine and harvested at AD14.

C) Quantification of *Bdnf* signal in medial VGLUT2 cells as *Bdnf* signal per cell per mouse injected with LacZ, P1, IE, or P4 gRNA AAV, treated with cocaine and harvested at AD14.

Statistics: n=4 mice, 50-70 cells sampled per animal, S3B: One-way ANOVA with post-hoc test being Tukey's multiple test, S3C: Nested One-way ANOVA with post-hoc test being Tukey's multiple test

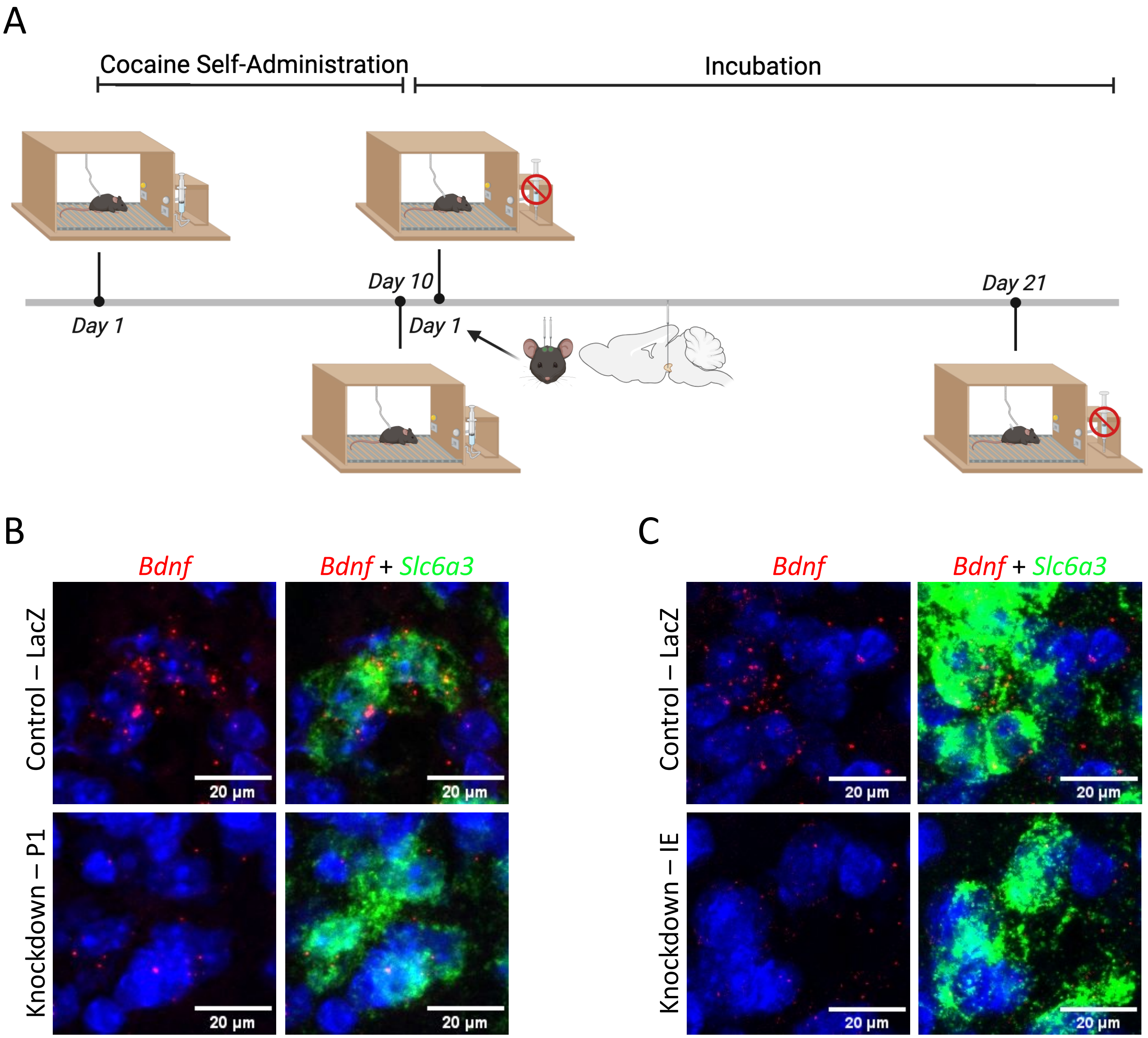


**Supplemental Figure S4: Validation of continued dCas9-KRAB-mediated repression of *Bdnf* after measuring incubation of cocaine craving behavior.**

A) Diagram representing incubation of cocaine craving paradigm

B) Representative RNA *in situ* hybridization images of *Bdnf* signal (red) in medial DAT cells (*Slc6a3* in green) from the VTA of mice injected with LacZ or P1 gRNA AAV

C) Representative RNA *in situ* hybridization images of *Bdnf* signal (red) in medial DAT cells (*Slc6a3* in green) from the VTA of mice injected with LacZ or IE gRNA AAV.

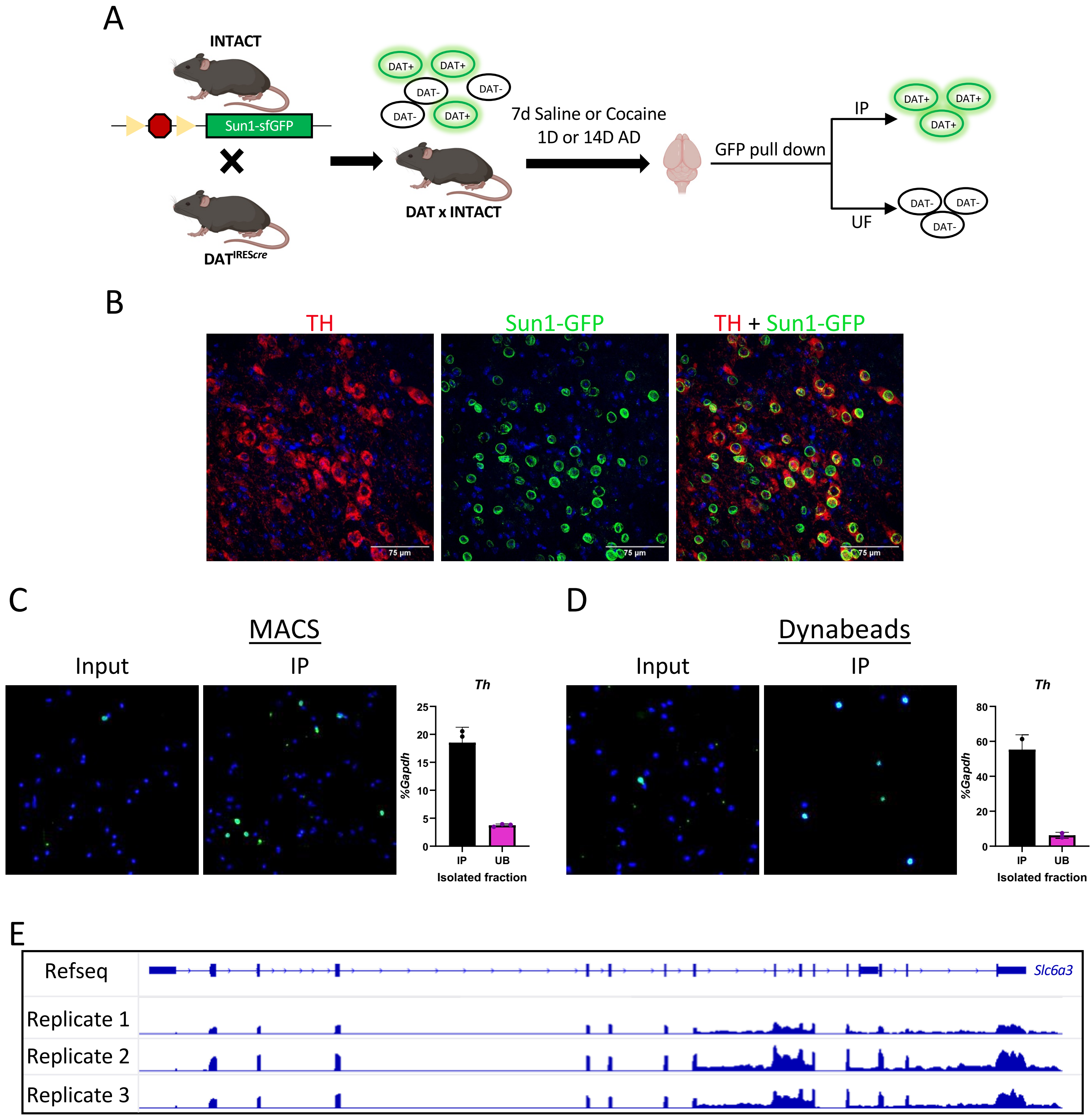


**Supplemental Figure S5: MACS and Dynabeads-dependent nuclei pull-down enriched DA nuclei from DATxINTACT mice**

A) Representative schematic of the generation of DATxINTACT mice and the enrichment of DAT+ nuclei

B) Representative immunohistochemistry image of tyrosine hydroxylase (TH, red) and nuclear membrane-bound GFP (Sun1-GFP, green) colocalization in the VTA of DATxINTACT mice

C) Representative image of DAT+ nuclei (green) before MACS pull down (left) and after MACS pull down (middle). RT-qPCR data showing enrichment of *Th* mRNA in the immunoprecipated (IP) fraction compared to the unbound (UB) fraction (right).

D) Representative image of DAT+ nuclei (green) before Dynabead pull down (left) and after Dynabead pull down (middle). RT-qPCR data showing enrichment of *Th* mRNA in the immunoprecipated (IP) fraction compared to the unbound (UB) fraction (right).

E) Representative igv tracks of RNA reads at *Slc6a3* from four samples that were enriched for DAT nuclei using Dynabead pull down.

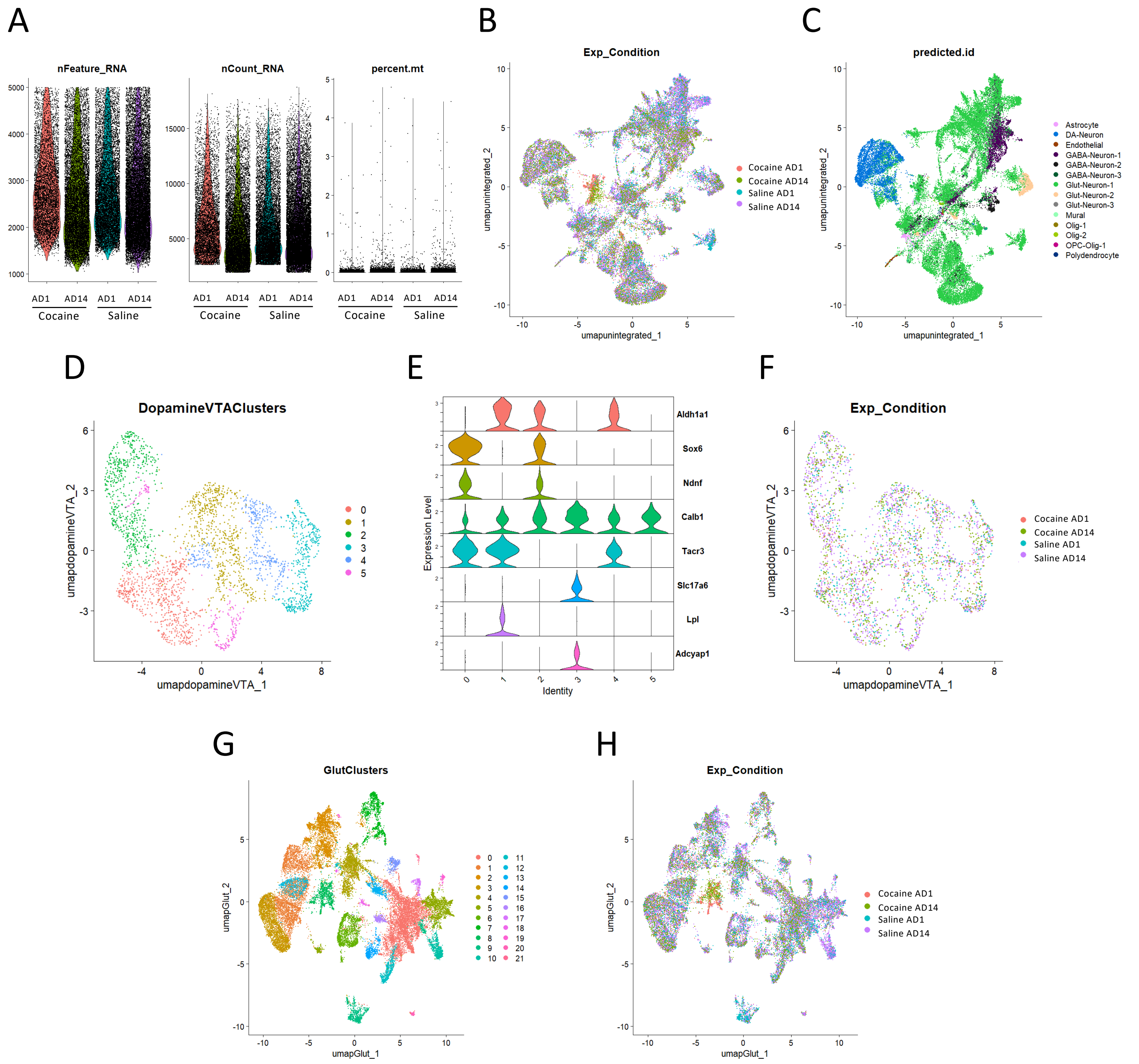


**Supplemental Figure S6: QC metrics and cell type identification in VTA snRNA-seq**

A) Distribution of snRNA-seq quality control metrics across conditions, including number of detected genes per cell (nFeature), total UMI counts per cell (nCount), and percentage of mitochondrial transcripts (percent.mt).
B) UMAP showing equal distribution of nuclei from all conditions across all VTA clusters
C) UMAP showing predicted cell types of each cluster, based on transferring anchors from a previously annotated VTA snRNAseq dataset.
D) Subclustering of the dopaminergic cluster and representation by UMAP reveals dopaminergic subpopulations.
E) Violin plots of key gene markers differentiating dopaminergic subpopulations.
F) UMAP showing equal distribution of nuclei from all conditions across all Dopamine clusters
G) UMAP of all glutamatergic nuclei from the total VTA snRNASeq dataset, with each color demarcating a cluster.
H) UMAP showing equal distribution of nuclei from all conditions across all Glutamatergic clusters.

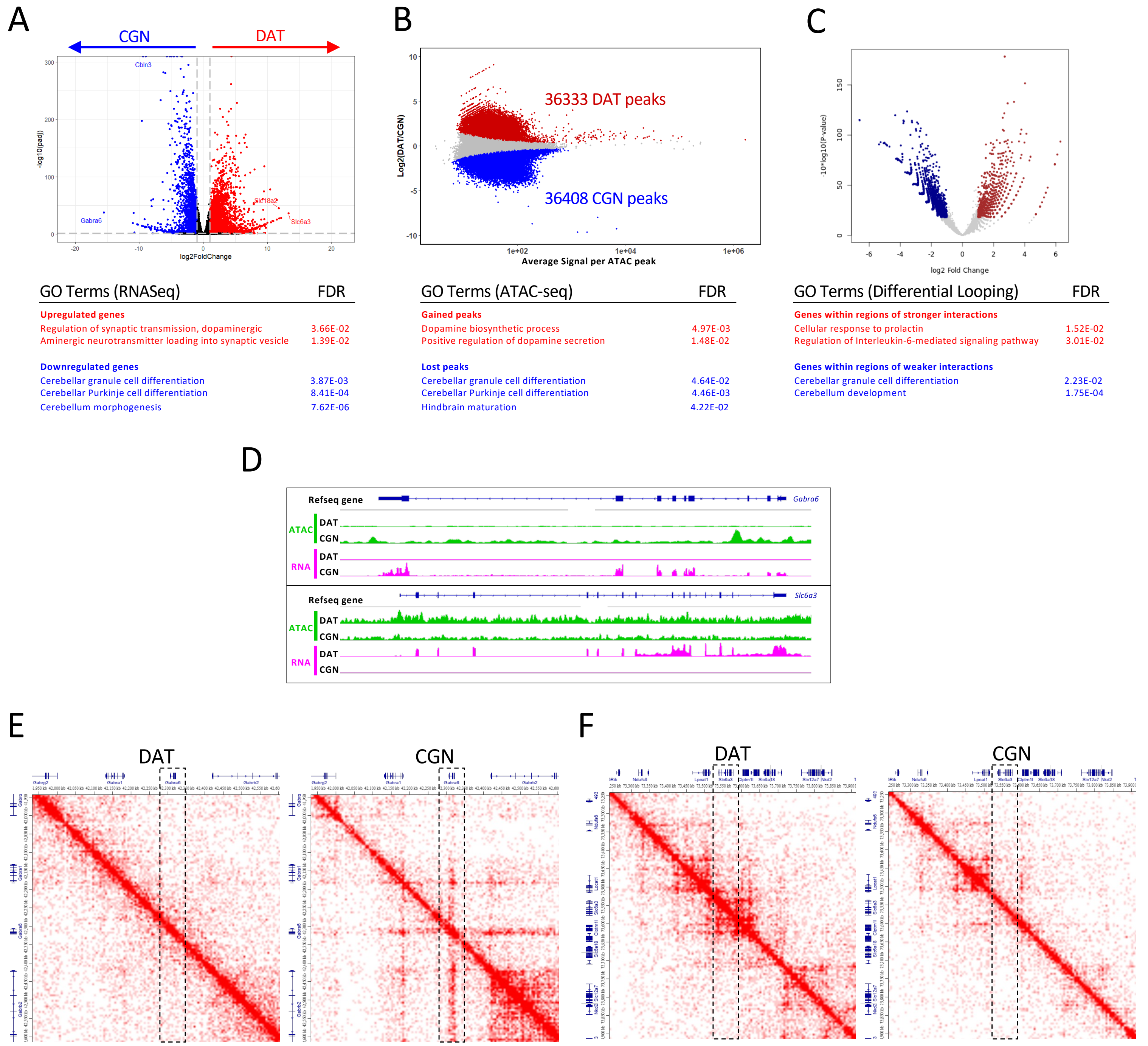


**Supplemental Figure S7: HiCAR can detect cell-type specific differences in chromatin accessibility and chromatin interactions**

A) Volcano plot (above) showing differential RNA expression when comparing DAT to CGNs, with upregulated genes enriched in DATs in red and downregulated genes enriched in CGNs in blue. GO terms for these differential genes (below)

B) MA plot (above) showing differential ATAC-seq sites when comparing DAT to CGNs, with gained ATAC sites (enriched in DATs) in red and lost ATAC sites (enriched in CGNs) in blue. GO terms for the genes possessing differential ATAC-seq sites at their promoters (below)

C) Volcano plot (above) showing differential chromatin interaction frequency when comparing DAT to CGNs, with increased interactions (enriched in DATs) in red and decreased interactions (enriched in CGNs) in blue. GO terms for the genes within these differentially interacting regions (below)

D) Representative ATAC-seq (green) and RNA-seq (magenta) tracks at *Gabra6* and *Slc6a3*, showing higher chromatin accessibility and gene expression of *Gabra6* in CGNs (top panel) and higher chromatin accessibility and gene expression of *Slc6a3* in DATs (bottom panel)

E) Juicebox images showing increased chromatin interactions at Gabra6 (dashed box) in CGNs compared to DATs

F) Juicebox images showing increased chromatin interactions at Slc6a3 (dashed box) in DATs compared to CGNs

Statistics: n=2-4 biological replicates, S7A-C: Differential enrichment calculated using DESeq2 package, FDR <0.05 and |L2FC|>1

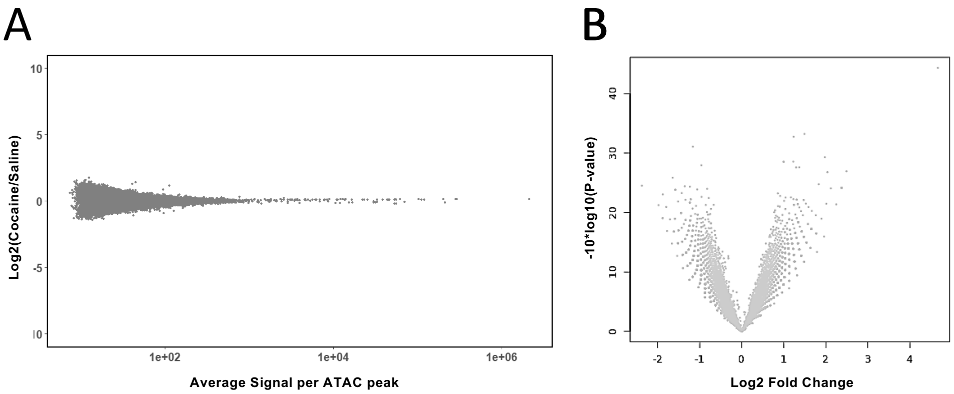


**Supplemental Figure S8: HiCAR analysis does not detect differences in chromatin accessibility and chromatin interactions in DAT cells between Saline AD14 and Cocaine AD14 conditions**

A) MA plot showing differential ATAC-seq sites when comparing Cocaine AD14 to Saline AD14
B) Volcano plot showing differential chromatin interaction frequency when comparing Cocaine AD14 to Saline AD14

Statistics: n=4 biological replicates per treatment condition, 2 male, 2 female, with 1 mouse/library. Differential enrichment calculated using DESeq2 package, FDR <0.05 and |L2FC|>1

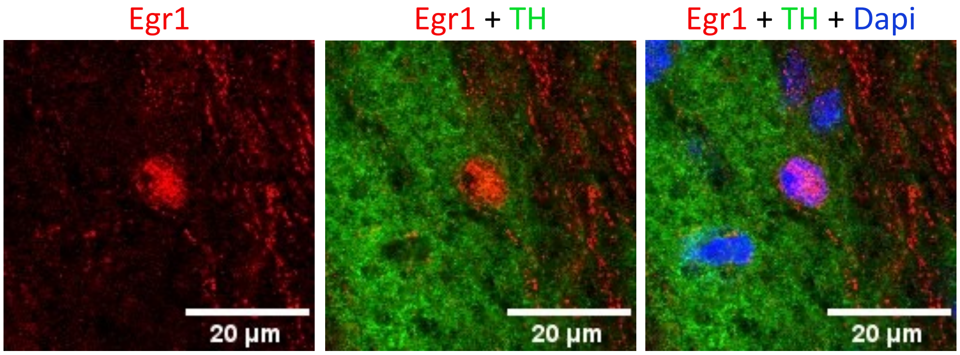


**Supplemental Figure S9: Egr1 positive cells are in regions of high TH expression.** Representative image of a Egr1 positive cell (red, left) colocalizing with high TH expression (green, middle). Merged image on the right with Dapi signal (blue).

**List of Supplemental Tables**

Supplemental Table S1: gRNA Sequences
Supplemental Table S2: Primer Sequences
Supplemental Table S3: scRNAseq Dopaminergic Cells MAST Results
Supplemental Table S4: scRNAseq Glutamatergic Cells MAST Results
Supplemental Table S5: GO Analysis Input Genes
Supplemental Table S6: HiCAR Primers
Supplemental Table S7: 1D ATAC-seq QC metrics

**Supplemental Table S1: gRNA sequences**

| gRNA | Sequence |
|---|---|
| LacZ | GCGAATACGCCCACGCGAT |
| P1g3 | ATGACTAGGCGAGAGGCACC |
| P1g5 | TCGCCTTGTCAAGCTAGGGC |
| P4g1 | TATATACTTAACCAGGATTT |
| P4g3 | ATTTCATGCTAGCTCGCCG |
| IEg3 | CAACTGGTTGCCGCTGGGTT |
| IEg5 | CCCAGCGGCAACCAGTTGGA |

**Supplemental Table S2: Primer Sequences**

| Primer | Sequence |
|---|---|
| RT-qPCR *Gapdh* F | 5’-CATGGCCTTCCGTGTTCCT-3’ |
| RT-qPCR *Gapdh* R | 5’-TGATGTCATCATACTTGGCAGGTT-3’ |
| RT-qPCR *Bdnf I* F | 5’-GCATCTGTTGGGGAGACAAG-3’ |
| RT-qPCR *Bdnf IV* F | 5’-CGCCATGCAATTTCCACTATCAATAA-3’ |
| RT-qPCR *Bdnf* R | 5’-GCCTTCATGCAACCGAAGTA-3’ |
| RT-qPCR *Pcsk1* F | 5’-TTGTTAATGAATGGGCGGCG-3’ |
| RT-qPCR *Pcsk1* R | 5’-CTCCGAGGATGGCTTTTGTG-3’ |
| RT-qPCR *Nr4a2* F | 5’-CCAATCCGGCAATGACCAG-3’ |
| RT-qPCR *Nr4a2* R | 5’-GATCTTCTCTGCCCACCCTC-3’ |
| HiCAR SplintV2 | 5’- TGTGCGAACTCAGACC-3' |

**Supplemental Table S6: HiCAR primers**

| Sample | I5 primer | I7 primer |
|---|---|---|
| CF1 | NEB primer i502 | Nextera-pcr-i7-2-L |
| CM1 | NEB primer i502 | Nextera-pcr-i7-3-L |
| SF1 | NEB primer i502 | Nextera-pcr-i7-4-L |
| SM1 | NEB primer i502 | Nextera-pcr-i7-5-L |
| CF2 | NEB primer i502 | Nextera-pcr-i7-6-L |
| CM2 | NEB primer i502 | Nextera-pcr-i7-7-L |
| SF2 | NEB primer i502 | Nextera-pcr-i7-10-L |
| SM2 | NEB primer i502 | Nextera-pcr-i7-11-L |

**Supplemental Table S7: 1D ATAC-seq QC metrics**

| Sample | TSSE Score | FRiP | bodyEnrich | promoterEnrich |
|---|---|---|---|---|
| DAT | 1.543918792 | 7.577870798 | 101672 | 116090 |
| CGN | 1.797455194 | 11.19841781 | 98657 | 118189 |
| Saline DAT | 1.486356542 | 9.432730972 | 102134 | 116328 |
| Cocaine DAT | 1.502673462 | 9.524374118 | 101769 | 116593 |
